# Microchromosomes are building blocks of bird, reptile and mammal chromosomes

**DOI:** 10.1101/2021.07.06.451394

**Authors:** Paul D. Waters, Hardip R. Patel, Aurora Ruiz-Herrera, Lucía Álvarez-González, Nicholas C. Lister, Oleg Simakov, Tariq Ezaz, Parwinder Kaur, Celine Frere, Frank Grützner, Arthur Georges, Jennifer A. Marshall Graves

## Abstract

Microchromosomes, once considered unimportant shreds of the chicken genome, are gene rich elements with a high GC content and few transposable elements. Their origin has been debated for decades. We used cytological and whole genome sequence comparisons, and chromosome conformation capture, to trace their origin and fate in genomes of reptiles, birds and mammals. We find that microchromosomes as well as macrochromosomes are highly conserved across birds, and share synteny with single small chromosomes of the chordate amphioxus, attesting to their origin as elements of an ancient animal genome. Turtles and squamates (snakes and lizards) share different subsets of ancestral microchromosomes, having independently lost microchromosomes by fusion with other microchromosomes or macrochromosomes. Patterns of fusions were quite different in different lineages.

Cytological observations show that microchromosomes in all lineages are spatially separated into a central compartment at interphase and during mitosis and meiosis. This reflects higher interaction between microchromosomes than with macrochromosomes, as observed by chromosome conformation capture, and suggests some functional coherence. In highly rearranged genomes fused microchromosomes retain most ancestral characteristics, but these may erode over evolutionary time; surprisingly *de novo* microchromosomes have rapidly adopted high interaction.

Some chromosomes of early branching monotreme mammals align to several bird microchromosomes, suggesting multiple microchromosome fusions in a mammalian ancestor. Subsequently multiple rearrangements fueled the extraordinary karyotypic diversity of therian mammals.

Thus microchromosomes, far from being aberrant genetic elements, represent fundamental building blocks of amniote chromosomes, and it is mammals, rather than reptiles, that are atypical.

**Significance Statement:** Genomes of birds and reptiles, but not mammals, consist of a few large chromosomes and many tiny microchromosomes. Once considered unimportant shreds of the genome, microchromosomes are gene rich and highly conserved among bird and reptiles, and share homology with one or more of the tiny chromosomes of an invertebrate that diverged from the vertebrate lineage 684 million years ago. Microchromosomes interact strongly and crowd together at the centre of cells, suggesting functional coherence. Many microchromosomes have been lost independently in turtles, snakes and lizards as they have fused with each other, or with larger chromosomes. In mammals they have completely disappeared, yet some chromosomes of the basal platypus line up with several microchromosomes, suggesting that they are the building blocks of the atypically variable chromosomes of mammals.

## Introduction

Classic cytological studies described mammalian chromosomes of a size easily visible under the microscope. Bird and reptile karyotypes were strikingly different, with an abrupt size discontinuity between macrochromosomes, with sizes (3-6 µm) in the range of mammalian chromosomes, and microchromosomes (<0.5 µm) which looked more like specks of dust (e.g. 1, 2, 3). These microchromosomes stained oddly and occupied a central position at mitosis (4).

An early view of microchromosomes as inconstant heterochromatic elements (5) or even not chromosomes at all, was thoroughly debunked (1, 6-8). Like macrochromosomes, they possess a centromere and telomeres at each end (with extra-long subtelomeric repeats) (9), and segregate regularly at mitosis. Microchromosomes are GC-rich and gene dense with a low content of repetitive sequence (10), and have high rates of recombination. They replicate early and are hyperacetylated compared to macrochromosomes, suggesting they are highly transcribed.

At the cytological level, most birds have extremely conserved karyotypes, including 9 pairs of macrochromosomes and 30-32 pairs of microchromosomes (3). Chromosome constitutions of birds are listed in (11). Though there are exceptions, even distantly related birds such as chicken an emu share nine macrochromosome pairs identified by banding patterns, chromosome painting and gene mapping (8, 12-14). Microchromosomes are too small to distinguish morphologically, let alone by G-band patterns, and pairing them is mostly guesswork. However, their number is usually constant and even, as expected for paired autosomes in diploids. Cytological examination, using specific DNA probes, suggests conservation of microchromosomes across 22 avian species (15), and comparative gene mapping and whole-genome analysis attests to considerable conservation among distantly related bird groups (16, 17). Genome sequencing of many bird species now provides unprecedented detail sufficient to compare microchromosomes across avian species.

Fewer comparative studies of microchromosome conservation have been done in non-avian reptiles, but their genome structures are similar to those of birds, with an abrupt distinction between a few macrochromosomes and many microchromosomes (reviewed in 18, 19, 20). However, turtles and snakes have fewer microchromosomes than birds. There is G-band and chromosome painting homology between the macrochromosomes of birds and turtles (21), and a close relationship between the chromosomes of birds and squamates (snakes and lizards) was noted early (22). Gene mapping, and sequence comparisons reveal many homologous synteny blocks (8, 23). Lizard karyotypes are more variable; some species have clearly demarcated macro and microchromosomes, whereas others show no clear distinction.

There are exceptional reptile and bird clades in which no abrupt size difference defines microchromosomes, and the size range of microchromosomes can also vary between clades. For example, eagle and parrot genomes have few microchromosomes (24) and crocodilians have 5 very large macrochromosomes and few chromosomes in the microchromosome size range (25).

The origin of microchromosomes has been debated for decades. Initially they were thought to represent some sort of breakdown product of “normal” mammalian-like macrochromosomes that existed in amniote and even tetrapod ancestors (26), and this view is still expressed (e.g. 27). The alternative view is that at least some of them represent the small chromosomes of a vertebrate ancestor 400 Mya, retained intact by several vertebrate clades (8, 28). Similarities with the small chromosomes of amphioxus (the lancelet, an early branching chordate) now suggest a much earlier origin (29), dating back to at least 684 Myr since they last shared a common ancestor with vertebrates.

With the availability of several chromosome-scale assemblies of bird and reptile genomes (10, 30), it is now possible to trace the origin and fate of microchromosomes in birds, reptiles and mammals. We compared the genomes of seven birds and ten reptiles with chromosome-level assemblies, as well as three mammals and an amphioxus (Figure 1). These comparisons provide evidence that, indeed, microchromosomes represent a set of highly conserved ancient animal chromosomes, whereas macrochromosomes, which are considered “normal” because of their ubiquity in mammals, have undergone multiple lineage-specific rearrangements, especially in mammals. We gather evidence that microchromosomes retain a high frequency of interchromosome interaction inside the nucleus, and regularly locate together at interphase and division, suggesting retention of an ancestral functional coherence between a set of small ancestral chromosomes.

**Figure 1.**
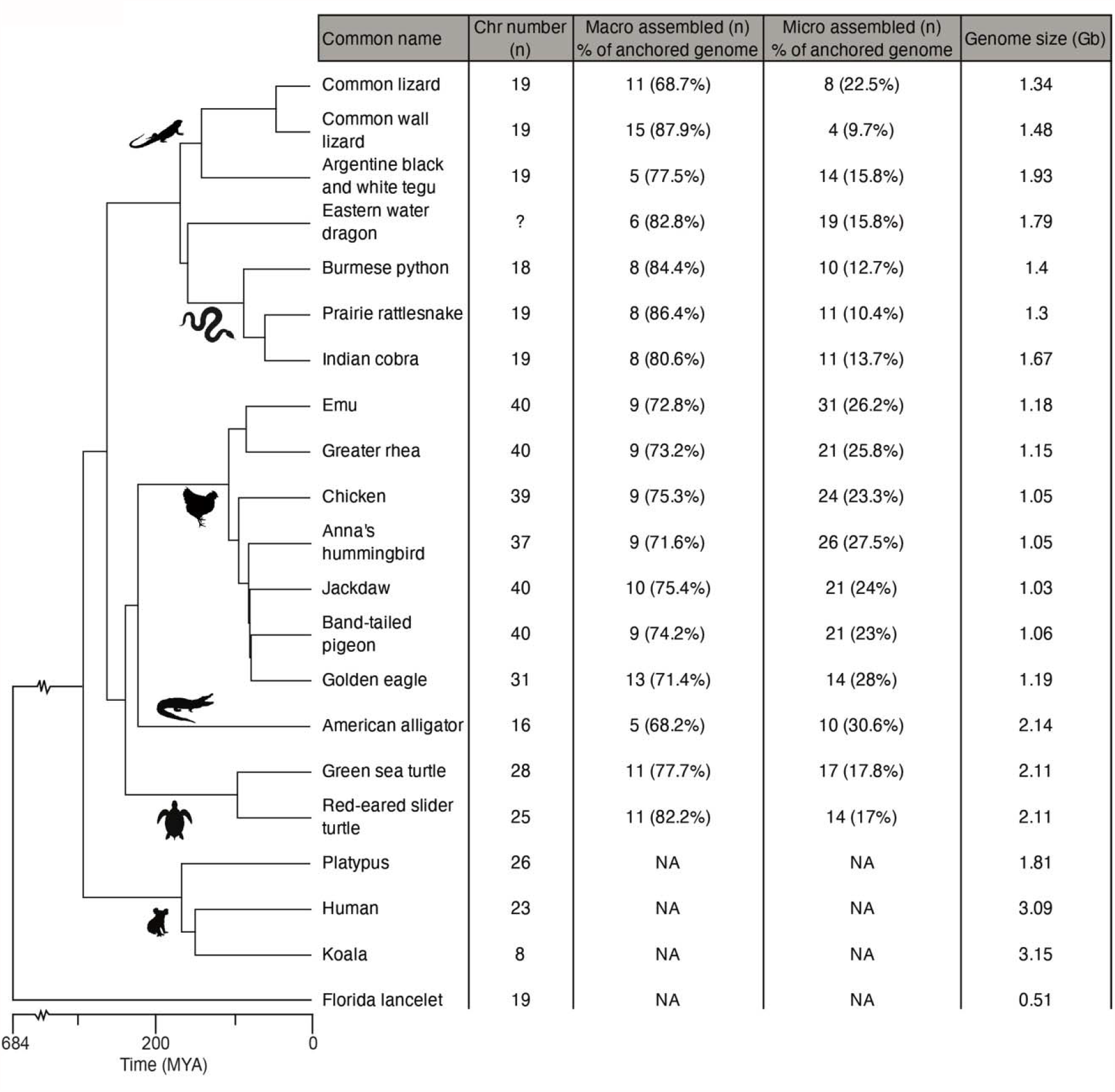
Phylogenetic relationships of reptiles, mammals and amphioxus genome assemblies compared in this study. Cytological chromosome numbers (n) are shown, along with the number of assembled macrochromosomes and microchromosomes (their percentage of the anchored genome), and genome size. Species names and full common names are given in Table S1: in the text they are referred to by abbreviated common names.

## Results

### Cytological observations of microchromosomes

To broaden our knowledge of microchromosome cytology, we examined the microchromosomes of several reptiles, including our model, the central bearded dragon *Pogona vitticeps*. In snakes (python and tiger snake), lizards (spiny-tailed monitor and bearded dragon) and a turtle (eastern long-neck) we found that microchromosomes were less strongly stained and tended to clump centrally in mitotic (Figure 2A) and meiotic cells (Figure 2B). Several repetitive sequences specifically hybridize to most or all the microchromosomes, allowing us to detect their central position within the interphase nucleus (Figure 2C).

**Figure 2.**
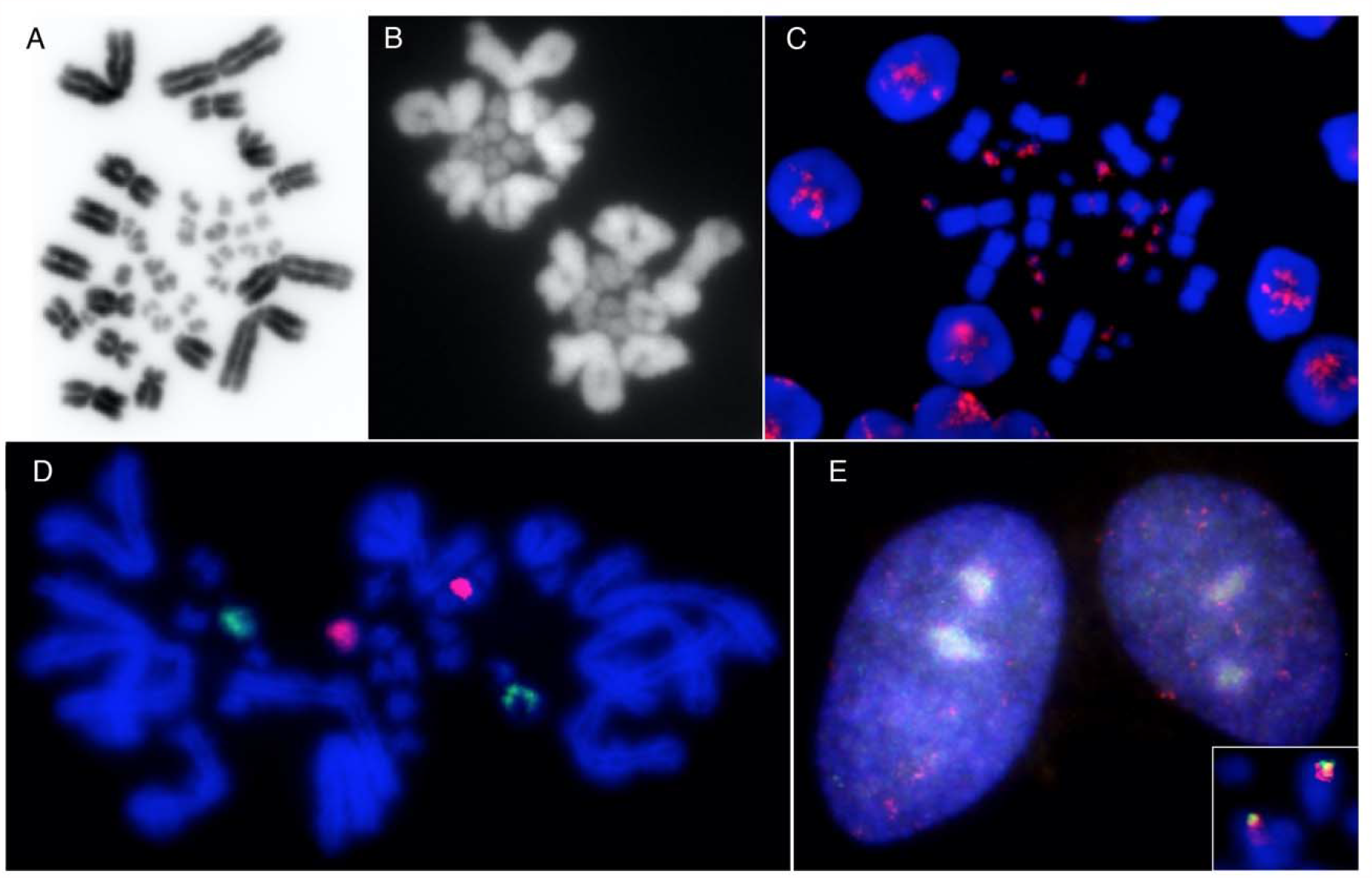
Cytological characterization of microchromosomes in reptiles. (A) Mitotic chromosomes of the bearded dragon, showing extreme size difference, different staining and central location of microchromosomes. (B) Polar view of diakinesis in bearded dragon spermatocyte, showing extreme size and staining difference and central location of macro and micro bivalents. (C) FISH using microchromosome-specific repetitive sequence [AGAT]_n_ showing central clustering of labelled dragon microchromosomes in interphase nuclei. (D) Two probes (green and red) light up two pairs of microchromosomes in dragon cells. (E) The same probes retain their central location in interphase nuclei although they co-locate to the terminus of a macrochromosome in the long-neck turtle (inset).

We also used DNA paints that specifically hybridized to two bearded dragon microchromosome pairs (Figure 2D) to explore their conservation in other species. We found that in the eastern long-neck turtle the two paints hybridized together at the tip of a macrochromosome, implying that these two sequences are fused, and fused to a macrochromosome in the turtle lineage; significantly, the paints hybridized to central regions of interphase nuclei in turtle as well as bearded dragon (Figure 2E).

Together our observations support the view that microchromosomes are differentiated from macrochromosomes, not only by their much smaller size, but also by their different staining properties (denoting different sequence make-up and chromosome conformation) and their location together in the centre of the interphase nucleus and dividing cells.

### Genome sequence comparisons

We performed pairwise whole genome alignments using LastZ (31) to identify syntenic blocks (reciprocally best aligned chains) and show conservation of chromosomes from birds, turtles and squamates within and between lineages. The relationships of the species we used are presented in Figure 1. We define microchromosome according to published karyotypes (see detailed information about species and their chromosomes in Table S1). In birds and snakes all assembled macrochromosomes were smaller than 35 Mb. In other clades the threshold size of the largest microchromsome was greater: i.e. lizards with rearranged genomes (50 Mb), tegu (75 Mb), alligator (96 Mb) and turtles (45 Mb).

### Sequence comparisons of bird and turtle genomes

Of the bird species with chromosome-scale assembled genomes, we chose emu, chicken, pigeon, jackdaw and hummingbird to capture the deepest avian divergences (Figure 1). Genomes of these five birds display striking homology, aligning >87% of their length (Figure 3A). The nine macrochromosomes (eight autosomes and the Z sex chromosome) are almost invariant. We observed no fusions of macrochromosomes, and only two macrochromosomes in hummingbird and one in jackdaw that have undergone fission. We conclude that macrochromosomes are extremely conserved within the avian clade.

**Figure 3.**
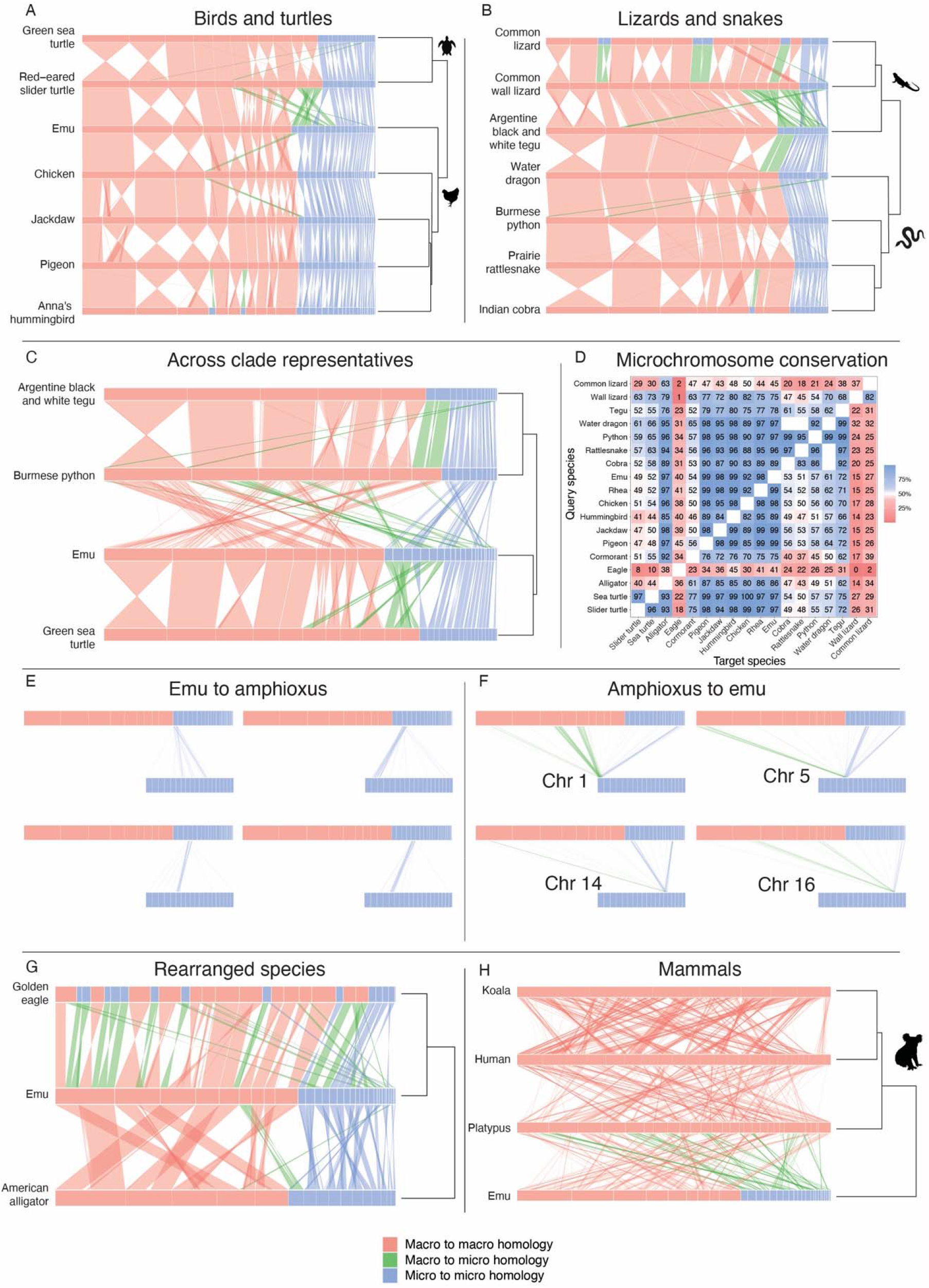
Sequence homology plots within and between birds, reptiles and mammals and comparison to the chordate amphioxus. Relationships between species, and genome sizes, are shown on the right. Genomes are resized, with chromosome sizes depicted as a proportion of genome length. Some chromosomes have been reordered in the plots to better show homologies within and between clades (see Table S2). Macrochromosomes are shown in red and microchromosomes in blue; changes between these states are shown in green. Chained and netted alignments were filtered for a minimum length of 100 kb for vertebrate species and 5 kb for amphioxus. Sequence homology between macro and microchromosomes within (A) birds and turtles, and (B) squamates. (C) Sequence homology between macro and microchromosomes of a representative lizard (tegu), snake (python), bird (emu) and the green sea turtle. (D) Heatmap showing the fraction (as a function of alignment chain length) of microchromosomes in the query species (y-axis) that align to microchromosomes in the target species (x-axis). E-F. Comparisons of emu and amphioxus chromosomes: single emu microchromosomes have homology to one (or two) amphioxus chromosomes (E), single amphioxus chromosomes detect strong homology to one (or more) emu microchromosomes, as well as to macrochromosome regions (F). (G) Sequence comparisons between emu and the rearranged alligator and eagle genomes. (H) Sequence comparisons between emu and mammals: eutherian (human), marsupial (koala) and monotreme (platypus).

There was little variation in the number of microchromosomes across these five birds, ranging from 28 pairs in hummingbird to 31 in emu (Figure 1, Table S1 and S2). Sequence comparisons of assembled microchromosomes show that they too are highly conserved, nearly all showing a 1:1 correspondence between all five bird species (Figure 3A). The most prominent exception is a microchromosome in all other bird lineages that aligns to chicken chromosome 4p, as previously noted (8, 12), and is significant because it also has homology to the conserved region of the mammalian X chromosome (32). Uniquely, the hummingbird genome contains two chromosomes in the microchromosome size range that are parts of macrochromosomes in other birds, implying an origin by fission of macrochromosomes. Conserved microchromosomes are GC-rich and gene dense in all species (mean of 40 genes/Mb compared to 17 genes/Mb on macrochromosomes in chicken) (Figure 4, Figure S1).

**Figure 4.**
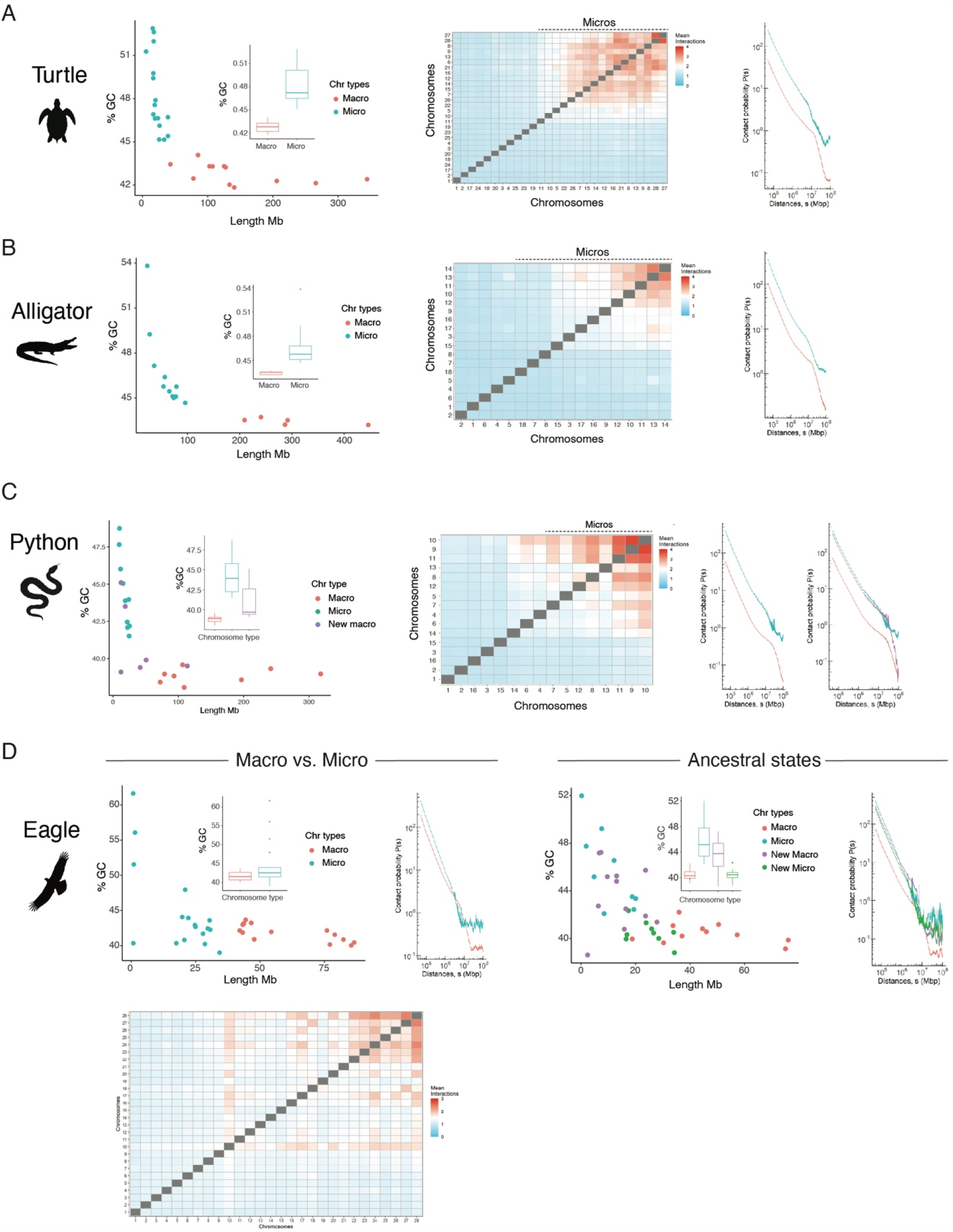
GC content versus size, inter-chromosome interaction ratios (with scaffolds sorted from largest to smallest, blue is low interaction and red high) and distance dependent interaction probabilities of macro and microchromosomes (Ps) for: (A) green sea turtle, (B) American alligator and (C) python. (D) GC content versus size, and distance dependent interaction probabilities of golden eagle macro and microchromosomes in the present derived state, and after partitioning the genome into ancestral micro and macrochromosomes and new micro and macrochromosomes

We conclude that microchromosomes and macrochromosomes are highly conserved between even the most distantly related bird lineages separated by ∼110 Myr.

We then broadened our comparison to include turtle genomes. Chromosome level assemblies of two turtle species were available – the green sea turtle and the red-eared slider turtle, with almost identical karyotypes. As in birds, turtle microchromosomes have a higher GC content than macrochromosomes (Figure 4, Figure S1). Sequence comparison shows 1:1 correspondence between two turtle genomes (Figure 3A) except for two green sea turtle microchromosomes that align to parts of large macrochromosomes in the slider.

Turtle macrochromosomes also align to emu macrochromosomes, though there are some intrachromosomal rearrangements. However, turtles have seven fewer microchromosomes. There is an apparent fusion of two microchromosomes conserved between sea turtle and emu that results in a larger microchromosome in the slider turtle. Four emu microchromosomes are present as two macrochromosomes in both turtle species. Two emu microchromosomes are fused to the termini of different macrochromosomes in sea turtle and slider, and one is fused to a macrochromosome in the sea turtle only.

Thus genomes in turtles, as well as bird, are highly conserved. Hereafter we use the emu genome (33) as a representative of the ancestral bird state and the green sea turtle to represent the turtle ancestral state, for inter-clade comparisons.

### Sequence comparisons of squamate genomes

Several genomes of snakes (python, rattlesnake and cobra) and lizards (common lizard, wall lizard, the distantly related tegu, and the outgroup water dragon) have been recently sequenced, and have chromosome-scale assembled microchromosomes to enable detailed comparisons and definition of ancestral states. The size differences between macro and microchromosomes were obvious in the snakes, water dragon and tegu, but less pronounced in the common and wall lizards. However, GC content and gene density of microchromosomes was higher than for the macrochromosomes in all these species (Figure 4A-C, Figure S1).

Sequence comparisons of snake genomes (Figure 3B) showed that macro and microchromosomes are almost wholly conserved between python, rattlesnake and the cobra. There were relatively few macrochromosome fusion/fission events between these species. The ten microchromosomes, too, were all conserved between the three snake species, apart from some minor rearrangements. However, notably, a cobra specific microchromosome aligns to part of a small macrochromosome in other species.

The water dragon, which is more closely related to snakes than the other lizards studied (34), shares the snake macro and microchromosome structure except for two microchromosomes that align to termini of the largest snake chromosome. Four water dragon microchromosomes had no alignments to any of the snake or lizard genomes. Chromosomes of the water dragon and tegu were almost identical (Figure 3B).

The common lizard and the wall lizard both had more and smaller macrochromosomes than tegu, water dragon or snakes; ten of these align to the five largest tegu and water dragon macrochromosomes, suggesting they are the products of fission. Both had few chromosomes in the microchromosome size range, and their alignments to multiple microchromosomes conserved between snakes, water dragon and tegu suggests fusion of two or three microchromosomes. There was also evidence of a triply fused microchromosome fused with another microchromosome in the wall lizard to form a chromosome out of the microchromosome range.

There is evidence, too, of microchromosomes arising *de novo* from chromosome fission. Four novel microchromosomes in the wall lizard have homology to two macrochromosomes in common lizard, which have homology to parts of two larger macrochromosomes in tegu. Another microchromosome shared by both lizards has homology to another region of the same tegu macrochromosome.

Thus, microchromosomes are very conserved among snakes, water dragon and tegu, and must represent the ancestral state. However, the wall lizard and common lizard lineages have undergone fusions (largely micro-micro) and a few fissions. We use the python and tegu to represent the ancestral state for lizards and snakes in across-clade comparisons.

### Sequence comparisons between birds and reptiles suggest that bird microchromosomes represent the ancestral amniote condition

We used pairwise genome alignments of the representative bird, turtle, snake and lizard for across-clade comparisons (Figure 3C). There was striking homology between the macrochromosomes, and also between the microchromosomes of emu, green turtle, python and tegu.

The genome sizes of turtles, snakes and lizards were larger than those of birds (Figure 3), suggesting that the sequence was expanded by the insertion and retention of repetitive elements. Comparisons of scaled genomes (Figure 3C) suggest that this expansion affected macrochromosomes and microchromosomes equally. Notably, a few emu microchromosomes expanded into the range of macrochromosomes in turtles or squamates; for instance, one microchromosome in tegu has 1:1 homology to a small chromosome in the common and wall lizards, but because it is larger than 50 Mb (our threshold for microchromosome size in squamates – see Table S1), it is classified as a macrochromosome in the wall (but not the common) lizard (Figure 3C).

Turtles have fewer microchromosomes than birds. Of the 22 assembled emu microchromosomes, only 15 are conserved as microchromosomes in the green turtle. A microchromosome present in the two turtle assemblies is missing from the emu and chicken assemblies. Four emu microchromosomes are present as micro-micro fusions. The other two emu microchromosomes align to arms of macrochromosomes in turtles, suggesting macro-micro fusion. Thus, turtles retain a subset of microchromosomes homologous to those of emu. Others are fused either with another microchromosome to form larger (micro or macrochromosome-scale) entities, or are fused to bird macrochromosomes.

Our representative squamates also have fewer microchromosomes than birds. Python has 10 and tegu 12 (plus two that align to a single macrochromosome in python and birds). The ten snake microchromosomes (and their tegu counterparts) all align to bird microchromosomes. Other bird microchromosomes align to autosomes in python and tegu, one to a terminal position, and seven others making up two chromosome arms. Thus squamates, too share a subset of bird microchromosomes.

Importantly, although nearly all of the microchromosomes in turtle and squamate assemblies are homologous to bird microchromosomes, they represent different subsets. Of 21 emu microchromosomes with sufficient homology to regions of both tegu and turtle genomes, only eight are microchromosomes in both (Figure 3C). Three are incorporated into different macrochromosomes in tegu and turtle. Seven emu microchromosomes have homology to turtle microchromosomes but squamate macrochromosomes, and three emu microchromosomes have homology to squamate microchromosomes but turtle macrochromosomes.

The simplest explanation of the pattern of microchromosome retention is that birds represent the ancestral amniote condition (31 microchromosome pairs), and micro-micro and micro-macro fusions reduced the numbers of microchromosomes independently in turtles and squamates.

Inspection of the bird microchromosomes with homology to macrochromosomes in turtles or squamates (or both) reveal different patterns of microchromosome fusion and fission. Among the bird microchromosomes with homology to regions of turtle macrochromosomes, there are fusions of two or three microchromosomes that generate larger (macrochromosome-sized) chromosomes. For instance, the two smallest turtle macrochromosomes are each homologous to two bird microchromosomes, implying fusions of ancestral microchromosomes. Other microchromosomes, or micro-micro fusions, have fused to terminal regions of macrochromosomes in turtles or squamates. Several appear to be Robertsonian (centric) fusions, in which an ancestral microchromosome or micro-micro fusion has become a macrochromosome arm (e.g. the two turtle macrochromosomes pictured in Figure 2E). Almost all other fusions are terminal, with very few examples of internal integration into a macrochromosome.

While the predominant evolutionary pattern is of continued loss of microchromosomes by fusion, a few novel microchromosomes have been created by fission from macrochromosomes, for example independently in hummingbird, cobra and common lizard.

To ensure that our inference about microchromosome conservation is not an artefact of selected pairwise comparisons, we calculated the percentage of microchromosome aligned regions in each query species, which aligned to microchromosomes of another target species (Figure 3D). We observed that 72-100% of microchromosome alignments for all species (except wall lizard and golden eagle) are to bird microchromosomes. When chicken (fusion of a microchromosome to chromosome 4) and hummingbird (fission of macrochromosomes that result in new microchromosomes) are excluded, >98% are micro-to-micro alignments between birds. It is intriguing to note that the heatmap (Figure 3D) is not symmetrical. This is because when birds are the target genome, most microchromosomes of other species align to bird microchromosomes. However, many of their microchromosomes have been fused into macrochromosomes in other clades, so when birds are the query genome the micro-to-micro alignment proportion is reduced. This demonstrates that the plot order of species in Figure 3 has no influence on our interpretation of microchromosome conservation and ancestral state.

### Origin of microchromosomes: homology with amphioxus chromosomes

The division of the bird genome into macro and microchromosomes represents the ancestral amniote condition, and the occurrence of microchromosomes in some fish (29, 35) suggests this characteristic might be ancestral to all vertebrates. But did microchromosomes arise by fission of larger chromosomes in an ancient chordate ancestor, or did larger vertebrate chromosomes arise by fusion of microchromosomes? Or did both processes occur (8)? To answer this question, we compared conserved reptile microchromosomes with the small chromosomes of a distant chordate relative, the amphioxus (Florida lancelet, *Branchiostoma floridae)*, which last shared a common ancestor with vertebrates 684 Mya.

Amphioxus has a small (520 Mb) genome divided into 19 tiny chromosomes that range in size from 17 Mb to 35 Mb. These chromosomes are very gene dense (60 genes/Mb compared to 10/Mb in mammals; Figure S1D). Comparison of the amphioxus sequence with those of garfish and chicken revealed two genome doublings; an autotetraploidization in the Cambrian ∼500 Mya and allotetraploidy by fusion of genomes that had diverged in a fish ancestor ∼460 Mya, followed by extensive loss of duplicate genes (29). Considerable sequence blocks shared synteny with the chicken genome, some of which represented 1:1 relationship with chicken microchromosomes. The Australian lungfish genome, though much expanded with repetitive sequence, also possesses many microchromosomes with homology to amphioxus chromosomes (35).

We aligned the bird (emu) and amphioxus genomes (Figure 3E, Figure S2). Of the 21 emu microchromosomes with sufficient shared sequence to detect homology, nine bird microchromosomes each represented a single amphioxus chromosome, suggesting that ancestral chromosomes have been retained intact. Another six emu microchromosomes each contained sequences from two amphioxus chromosomes, implying fusions in ancestral vertebrates.

We then aligned single amphioxus chromosomes to the emu genome, demonstrating that most have strong homology to one or two microchromosomes, as well as to two (sometimes three) regions of macrochromosomes (Figure 3F, Figure S3). These probably represent the four copies of the chordate genome that underwent two rounds of doubling. It is notable that one or two of these homologies (usually including at least one microchromosome) are strong and focused, whereas the others are more dispersed, suggesting rearrangement, deletion and repeat expansion. This suggests that all copies of each ancient chromosome have retained synteny (physical linkage), with a single copy equating to one or more microchromosomes as previously observed (29).

Unsurprisingly, given their homology to bird microchromosomes, most turtle, snake and lizard microchromosomes also equated to single or fused amphioxus chromosomes. The most parsimonious explanation is that reptile/bird microchromosomes represent ancestral chordate chromosomes (or fused chromosomes) that have retained synteny.

We conclude that reptile and bird microchromosomes are relics of an original animal genome with tiny, gene-rich chromosomes, represented today by amphioxus. Since turtle and squamate microchromosomes are different subsets of bird microchromosomes, this implies that these ancestral microchromosomes have been progressively lost by fusion at different rates independently in different lineages. Fusion has evidently been slower in birds than in other reptile lineages.

### Fusion of microchromosomes into macrochromosomes

To examine the process by which ancient microchromosomes became incorporated into macrochromosomes in vertebrates, we examined the genomes of exceptional reptile and bird species in which fusion has removed many or most microchromosomes. We aligned the genome of emu (representing ancestral birds) with alligator (representing crocodilians that diverged from birds 240 Mya), and eagle (which last shared a common ancestor with other birds with a conserved karyotype about 80 Mya).

We found that emu macro and microchromosome sequence had obvious homologues in the alligator and eagle genomes, although they were considerably rearranged (Figure 3G). Strong homology between emu and alligator macrochromosome arms show that the five very large alligator chromosomes are all fusions and rearrangements of ancestral macrochromosomes, as was demonstrated by chromosome painting between the chicken and the Nile crocodile (19, 36) which shares many chromosomes with alligator (25). An alligator macrochromosome and the largest microchromosome were each generated by fusions of an ancestral macro and microchromosome. However, the ten smallest alligator chromosomes each comprise either single or fused ancestral microchromosomes; one small alligator chromosome represents a single microchromosome, five alligator chromosomes represent fusions of two, and four alligator chromosomes fusions of three ancestral microchromosomes. These changes must have occurred in the crocodilian lineage in the ∼240 Myr since they shared a common ancestor with birds. It is striking that all but two rearrangements are either a macro-macro or a micro-micro fusion.

The rearranged eagle genome, in contrast, contains many fusions between macro and microchromosomes, and many fissions of ancestral macrochromosomes into smaller chromosomes (Figure 3G). Only three macrochromosomes have been retained intact, two have undergone centric rearrangements of chromosome arms and two have each fused with an ancestral microchromosome. The other four ancestral macrochromosomes have undergone multiple fissions, the products of which have fused to other macrochromosome arms or ancestral microchromosomes. The largest emu chromosome has undergone fission into seven regions, six of which are present as small chromosomes (five in the micro range). In addition, two small eagle macrochromosomes derive from micro-micro fusions.

Thus eagle microchromosomes include only four ancestral microchromosomes and one micro-micro fusion. Three ancestral microchromosomes have fused to form macrochromosome arms or terminal regions. What is striking is that six eagle microchromosomes were derived from regions (not all terminal) of ancestral macrochromosomes, so represent *de novo* microchromosomes.

Thus, in both alligator and eagle, macrochromosome arms and microchromosomes have been fused with each other, and with the termini of macrochromosomes. However, the patterns of fusions and fissions are quite different in the eagle and alligator, attesting to independent rearrangement facilitated by different mechanisms.

### Microchromosome interactions

Our cytogenetic studies (above – Figure 2) confirm and extend previous observations (4) that microchromosomes are spatially segregated within interphase and dividing cells, occupying a central location at interphase, and during mitosis and meiosis, in turtles and squamates as well as birds. We also showed that microchromosomes may retain this central position even after fusion to macrochromosomes, implying that size alone does not determine microchromosome location.

Data from high-throughput chromosome conformation capture (HiC) now provide a molecular description of this spatial segregation. HiC data incorporated in the new emu assembly (33) and the rattlesnake assembly (37) reveal that microchromosomes interact with each other more than with macrochromosomes in these species, as has been observed also in other birds, snakes and turtles (38). We confirmed and extended these observations to other birds and reptiles (green sea turtle, alligator, python, eagle, tegu, emu, greater rhea and water dragon), using DNA Zoo HiC data (39) (Figure 4A-B, Figure S1). These species show various degrees of rearrangement of ancestral macro and microchromosomes, permitting us to determine whether microchromosome fusion or macrochromosome fission can alter the GC ratio and interaction characteristics.

On average, pairs of loci borne on the same chromosome present higher interaction probabilities (intra-chromosomal interactions) than between loci on different chromosomes (inter-chromosomal interactions). At the genome-wide level, this cis/trans interaction pattern reflects chromosomal territoriality (40). As expected, the interaction between neighboring loci on the same chromosome decreases as genomic distance increases. This reduction in interactions can be represented as a genomic distance-dependent contact probabilities [P(s)], representing the level of chromosome compaction (41).

Genome-wide heatmap plots of representative species (Figure 4, Figure S1) all show that, as well as a high GC ratio, there is a higher interchromosomal interaction and higher distant dependent contact probabilities [P(s)] between microchromosomes than between macrochromosomes or between macro and microchromosomes. More intense interaction is therefore an intrinsic feature of all reptile/bird microchromosomes, reflected by their arrangement within the nucleus during interphase (Figure 2).

Given that this interaction pattern is an ancestral feature of microchromosomes, we asked whether rearrangements in alligator, water dragon, python and eagle resulted in changes of properties of ancestral and *de novo* microchromosomes.

The alligator has small chromosomes formed by micro-micro fusions (Figure 3G). We found that these fused microchromosomes retain their higher GC ratio and still interact strongly with each other and the smaller microchromosomes (Figure 4B).

In the water dragon, several ancestral microchromosomes have fused to form four new regions of macrochromosomes. These are also present in the python genome (Figure S1E), so must have occurred in a common ancestor about 180 Mya. When we plotted GC content and contact probabilities according to the ancestral state (Figure 4C, Figure S1F), we found that these ancestral microchromosomes fused to macrochromosomes had reduced GC content, but retained their high contact probability. However, two ancestral microchromosomes that were more recently fused to either end of the largest scaffold only in python still retain their elevated GC content (Figure 4). This suggests that the GC-richness of microchromosomes is retained on rearrangement, but erodes over time.

The much rearranged eagle genome, for which there is both chromosome level assembly and HiC data, allowed us to assess features of both microchromosomes which fused to form macrochromosomes, and macrochromosomes that were split into microchromosomes, in the ∼80 Myr since the eagle last shared a common ancestor with condors, which retain an ancestral bird karyotype (42).

Of the 30 ancestral microchromosomes present in other birds, only four remain in the eagle; two have fused into a larger microchromosome and 12 are fused with macrochromosomes. Nine new microchromosomes represent regions of ancestral macrochromosomes which underwent fission (Figure 3G).

Unlike other bird species (Figure S1), the eagle micro and macrochromosomes showed no abrupt differences in GC content or contact probabilities with size (Figure 4D), reflecting their recent reshuffling. The microchromosomes still retained elevated interchromosomal interactions (Figure 4D), although this was not as well correlated with chromosome size as for other species.

To explore the origin of these differences, we examined the characteristics of eagle sequence according to ancestral state as deduced from comparison with the emu genome (Figure 3G). Regions of synteny with the emu genome were classified as macro (macro in both species), micro (micro in both species), new macro (micro in emu but macro in eagle) or new micro (macro in emu but micro in eagle). We found that GC content reflected the ancestral state (Figure 4D). The microchromosomes incorporated into macrochromosomes (new macros) in eagle maintained high GC content, and macrochromosomes broken down to microchromosome size (new micros) retained their low GC content.

In contrast, the distance dependent contact probabilities [P(s)] of these four classes of regions did not consistently reflect their ancestral state. As expected, ancestral macrochromosomes displayed lesser, and microchromosomes greater distance dependent interaction probabilities (Figure 4D). Microchromosomes incorporated into macrochromosomes also maintained relatively high contact probabilities at genomic distances below 1 Mbp, albeit slightly lower than the ancestral microchromosomes. Surprisingly, however, the new microchromosomes derived recently from macrochromosomes had almost the same elevated distance dependent interaction probabilities as the ancestral microchromosomes. This cannot be a consequence of altered base ratio, since %GC was not increased, and may reflect different levels of chromatin compaction or compartmentalisation.

We conclude that the high interaction probabilities of ancestral microchromosomes are maintained after their incorporation into a macrochromosome, but erode over time. However, novel microchromosomes rapidly adopt high interaction with other microchromosomes.

### The fate of microchromosomes in mammals

To explore the fate of ancient microchromosomes in therian mammals (eutherian and marsupial mammals), we compared the genomes of emu with those of koala (a marsupial) and human (eutherian) (Figure 3H). We observed regions with some homology to bird microchromosomes, though these were weak and dispersed. There was one region of the human genome (chromosome 17) and two regions of the koala genome that had homology to two or more microchromosomes, but no evidence that microchromosomes are retained intact in either species.

Monotremes (egg-laying mammals such as the platypus) have a karyotype somewhat resembling those of reptiles, with 6 pairs of large chromosomes, and 20 pairs of much smaller chromosomes (43). Although not in the same size range as reptile microchromosomes, we asked whether these were the vestiges of ancestral microchromosomes. We found that most platypus chromosomes, large and small, had contributions from multiple chicken macro and microchromosomes, and there was no obvious enrichment of microchromosomes in the small platypus chromosomes (Figure 3H).

However, homology with ancestral microchromosomes was not distributed randomly within the platypus genome. Four platypus chromosomes appear to be entirely composed of regions with homology to several emu microchromosomes (4, 3, 2 and 1 respectively), suggesting that they evolved from multiple microchromosome fusions. Other regions with strong homology lay at the ends, or comprised arms of platypus chromosomes.

This suggests that in the reptile-like ancestor of mammals, micro-micro fusions may have been common (as they were also in the crocodilian ancestor). These blocks of fused microchromosomes have been preserved in platypus macrochromosomes, but were disrupted in therian mammals, so that little vestige remains. The high rate of genome reshuffling in mammals contrasts with the stability of genomes in other amniote lineages.

## Discussion

Our analysis of sequence data confirms that reptile and bird microchromosomes and macrochromosomes are highly conserved within bird, turtle and squamate lineages, and even between these lineages. Microchromosomes are most numerous and almost invariant among birds, many species having 31 well conserved microchromosomes. Different microchromosome subsets have been retained in turtles and squamates, implying that the 31 microchromosomes were present in the common ancestor of birds and reptiles about 300 Mya, expanding on the 20 considered by Burt et al (8) to be ancestral. A few bird species have more microchromosomes, but chromosome painting reveals their recent origin from fission (14).

Strong homology with the small chromosomes of amphioxus implies that the conserved set of bird microchromosomes represent retention of ancient chromosomes of a common ancestor that lived 684 Mya. Comparisons of single amphioxus chromosomes with the bird genome reveal homology to four regions, one or two with bird microchromosomes and two or three with bird macrochromosomes. Presumably these four regions of demonstrable homology reflect paralogous sequences generated by the two genome doublings early in vertebrate evolution (29, 44). For most of these paralogous regions, one or two of the strongest are located on bird microchromosomes. The strongest signals (many on microchromosomes) probably represent paralogous sequences that retain the gene richness and low repetitive sequence content of the original chromosomes, whereas other genome copies have suffered deletion, transposable element invasion and rearrangement. It is interesting to speculate that microchromosomes may be protected from rearrangement and insertion of repetitive elements by their longer subtelomere regions, their spatial isolation and high interaction.

Microchromosomes, as well as macrochromosomes have been proportionately lengthened by insertion of transposable elements, as observed over many vertebrates (45), occasionally moving out of the microchromosome range. However, most microchromosomes have been progressively lost by fusion, and very occasionally gained by fission of macrochromosomes, in all bird and reptile lineages (Figure 5).

**Figure 5.**
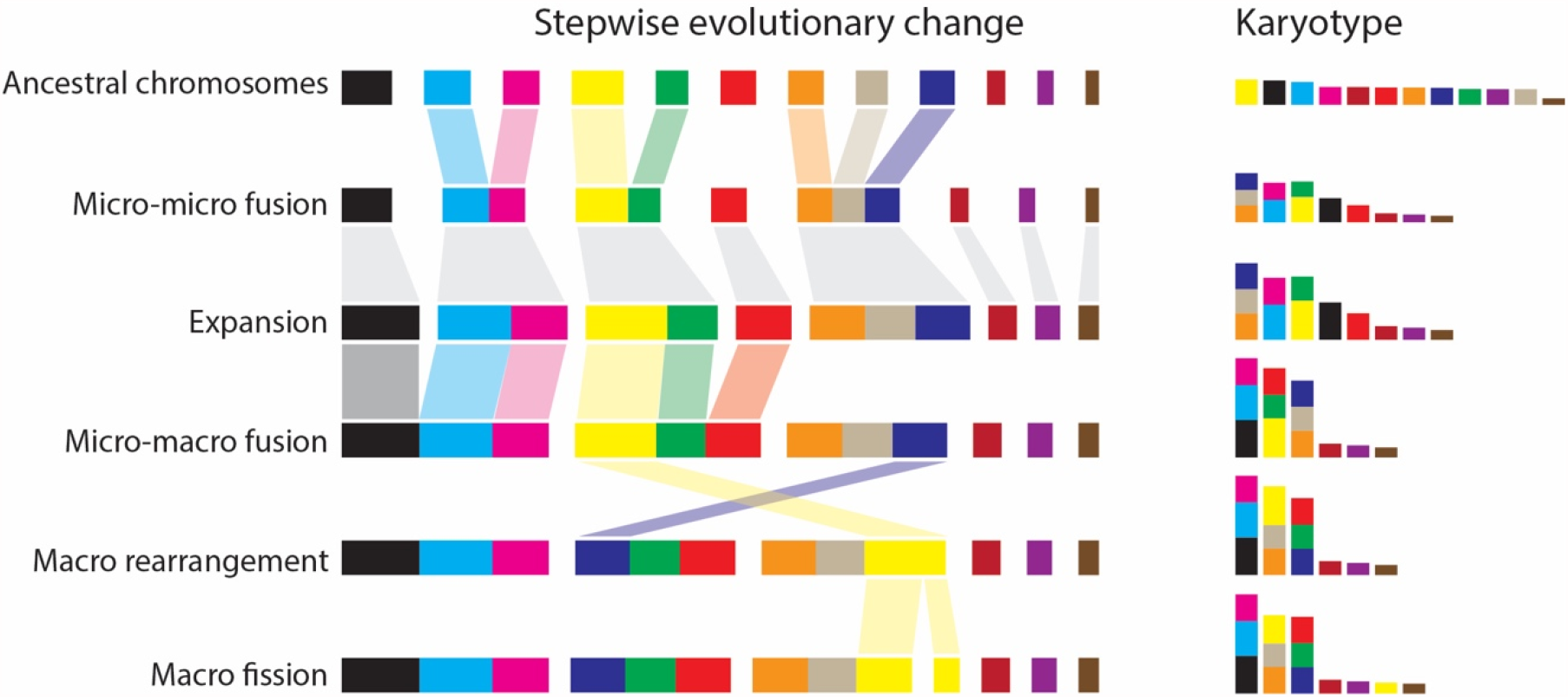
Evolutionary loss and gain of microchromosomes has reshaped the amniote kartyotype. Fusion between microchromosomes, expansion by the insertion of transposable elements, fusion with macrochromosomes and macrochromosome rearrangement has led to a reduction of microchromosomes. There are few macrochromosome fissions that result in new microchromosomes.

Patterns of microchromosome loss (Figure 5) were examined by comparing the genomes of conserved representatives of birds, turtles and squamates. Almost all loss can be attributed to fusion of two, or sometimes three, microchromosomes to form larger chromosomes, as well as fusion of microchromosomes to macrochromosomes. These fusions are almost always to terminal locations or via centric fusion to constitute a chromosome arm, as was implied by early observations of interstitial and centromeric telomere sequence (46).

Patterns of microchromosome loss were also examined by following the chromosome changes in exceptional species (alligator and golden eagle) with more rearranged chromosomes. We found that these genomes differ from the standard simply by more fusions of the same types that distinguish turtle and squamate genomes, suggesting that the same processes occurred, but at much higher rates.

The onset of instability in a lineage could be quite sudden. For instance, extensive genome remodeling in the eagle lineage must have occurred in the last 80 Myr since eagles shared a common ancestor with condors, which retain a near-ancestral genome arrangement (42). Cormorants diverged from the same conserved lineage a short time later. Our analysis of the cormorant assembly (47) (alignments not shown) revealed a rearranged genome, but few of the rearrangements are shared with eagles. The high rate of independent changes in cormorants or eagles suggests that instability was introduced to the genome of a common ancestor, but was expressed independently in the two lineages. One explanation of the sudden onset of instability after millions of years of extreme conservation is that the genome of an eagle ancestor (∼80 Mya) was invaded by a transposable element (TE), which facilitated interchromosome rearrangements (48).

Intriguingly, the patterns of microchromosome fusions are quite different in the two exceptional species we examined. Rearrangements in the alligator are almost all confined to micro-micro and macro-macro fusions, whereas in the eagle, they are largely micro-macro fusions. Assortative fusions (micro-micro and macro-macro) may be favoured by the spatial compartmentalization (49, 50). However, it is likely that rearrangements are driven by the location and type of repetitive elements inserted, as seems to be the case for parrots (24). Transposable elements are ubiquitous in reptiles and birds (51) but not randomly distributed in the genome (52). For instance, there are several repetitive sequences that are shared only by microchromosomes (53) (e.g. Figure 2C), which may favor micro-micro rearrangements, such as those that predominate in alligator. Other transposable elements are observed to cluster at centromeres of all chromosomes (54), and may be involved in Robertsonian fusions between macro and microchromosomes.

Loss of microchromosomes has, rarely, been offset in some lineages by the creation of novel microchromosomes from macrochromosome fragments. Several appear in the eagle genome, and there are one or two in other birds (e.g. hummungbird) and squamates (e.g. common lizard).

Reptile and bird microchromosomes are distinguished by their high gene density and high GC content, and particularly by their compartmentalisation in the centre of interphase and dividing cells. We have documented differential staining (denoting differences in sequence and conformation) and spatial segregation in a variety of squamates and turtles as well as birds, consistent with many cytological observations (4). The molecular underpinnings of these properties were revealed by our analysis of chromosome conformation capture data, which shows that microchromosomes in bird, turtle and squamate lineages all show high interaction within this compartment. These intense interchromosome relationships have been thought to denote different chromatin organization of microchromosomes that reflects some functional coherence.

We analysed rearranged genomes to examine properties of regions whose status has changed between micro and macrochromosome. We found that high interaction, as well as high GC content, is retained by ancestral microchromosomes that fused with macrochromosomes in eagle. This is consistent with our cytological observation that two ancestral microchromosomes fused to a macrochromosome in a turtle retain their central position (Figure 2E), as do fused microchromosomes in the rearranged genomes of falcons and parrots (55). However, high GC content does appear to erode with time, as shown by a lower GC content of fused microchromosomes in snakes.

We discovered, unexpectedly, that novel microchromosomes derived by fission of ancestral macrochromosomes also had an increased distance dependent interaction probability, although their GC content stayed low. This suggests that intense interaction may be the result, not of sequence composition, but of chromosome compaction or location in the cell, which may be influenced by epigenetic factors or chromosome size.

The extreme conservatism of the reptile genome, and even the modest rearrangements that characterize the crocodilians, expose the flagrant rearrangements in mammals as a glaring exception among amniotes. Mammals have extraordinarily variable karyotypes. In eutherian mammals, a near-identical-sized (3 Gb) genome is packaged as anything between 3 pairs of enormous chromosomes in the Indian muntjac to 51 pairs in the red viscacha rat (56). Not only is the highly rearranged eutherian genome subdivided into a few large or many small chromosomes, but sequences have been scrambled in many lineages. Large synteny blocks are shared by some species (e.g. humans, cats and even sloths), enabling reconstruction of ancestral eutherian karyotypes (57, 58), but even these blocks of shared synteny are wildly different in other species, especially the rodents. The marsupial karyotype, in contrast, is highly conserved between all 260 species, and derives from a n=7 basal karyotype largely by Robertsonian translocations (59).

We found that in eutherians and marsupials, microchromosomes have completely disappeared, visible only as broken up patches of homology peppering the genome.

Even the early branching monotreme mammals, which have a rather reptilian karyotype with 6 large and many small chromosomes, retain none of the ancestral microchromosomes. Most of the small platypus chromosomes have homology to several regions of macro and microchromosomes. However, two large and two small platypus chromosomes seem to be completely made up of fusions of several ancestral microchromosomes, suggesting that the process of amalgamation may have started from many micro-micro fusions.

We propose that microchromosome fusion occurred in the ancestor of all mammals after divergence from the reptile/bird lineage 310 Mya and before the divergence of monotremes from therians 188 Mya. These microchromosome blocks must have already undergone major sequence reshuffling before the marsupial-eutherian divergence 168 Mya because microchromosome-homologous sequences are split up and distributed all over the genome in both lineages.

The sequence shuffling and size variation among eutherian chromosomes would require some event that loosened the constraints on rearrangement in a mammalian ancestor, probably invasion and amplification of particular retrotransposons, which provide sites for crossing over between non-homologues (reviewed 60).

Thus, mammal genomes are spectacularly atypical among amniotes, displaying variation that has been sometimes credited with their rapid speciation and success (e.g. 61). We need to understand what effects these rearrangements between ancestral macro and microchromosomes had on genome function. The high gene density, atypical base ratio, spatial segregation, and high interaction between microchromosomes suggests a functional coherence of this part of the genome, which survives in the subsets of microchromosomes retained in birds, turtles and squamates. Gene-dense and active chromosome regions are also located centrally in mammalian cells (62), but these do not equate with ancestral microchromosomes. It is difficult to unscramble cause and effect relationships between chromosome size, location, gene density, GC content, and distance dependent interaction probabilities.

Our overall conclusion is that bird microchromosomes represent remnants of the original building blocks of vertebrate genome. They retain high gene density and low content of repetitive sequence, and share conserved features across all reptile and bird clades. Their progressive fusion with each other, and with macrochromosomes, occurred conservatively and gradually in most reptile lineages, but more rapidly in a few clades. However, multiple microchromosome fusions occurred early in mammal evolution, and was followed by lineage-specific rearrangement and a huge variety of fusions and fissions that disrupted the relationship between microchromosomes. Among amniotes, even vertebrates, mammal genomes are the true exceptions.

## Materials and Method

### Cytology

Mitotic and meiotic chromosomes preparations, chromosome paint preparation and painting were performed following protocol described in (63). Repeat mapping was performed following protocol described in (64). Briefly, 200 ng 5’-Cy3-labeled (AAGG)_8_ probe was mixed with 15 μl hybridisation buffer (50% formamide, 10% dextran sulfate, 2× SSC, 40 mmol/L sodium phosphate pH 7.0 and 1× Denhardt’s solution) added to slides containing fixed chromosome preparation and denatured at 68°C for 5 min. After denaturation, slides were incubated overnight in a moist hybridisation chamber at 37°C. Slides were then washed once at 60°C in 0.4× SSC, 0.3% Igepal for 2 min, followed by another wash at room temperature in 2× SSC, 0.1% Igepal for 1 min and air-dried. Slides were then counter stained with DAPI with Vectashield. Image analysis was performed using a Zeiss Axioplan epifluorescence microscope equipped with a CCD (charge-coupled device) camera (RT-Spot), (Zeiss, Oberkochen, Germany).

### Whole genome alignments

One way all-versus-all LastZ (Release 1.02) (31) alignments were performed for 23 reptile (including birds) species and platypus. Human, Tasmanian devil and amphioxus genomes were aligned against selected genomes. Alignments were chained and netted using the UCSC Toolkit (http://hgdownload.soe.ucsc.edu/admin/exe/linux.x86_64). Workflow was automated using scripts available at https://github.com/kango2/tiny.

Briefly, the LastZ alignment parameters were: K=2400 L=3000 Y=9400 H=2000 -- ambiguous=iupac. Chaining was performed with axtChain using -minScore=3000 - linearGap=medium as parameters. Chains were sorted with chainSort, pre-netting was performed with chainPreNet, and netting performed with chainNet, each using default parameters. Syntenic blocks were calculated with netSyntenic.

Homology and statistics were plotted in R using the tidyverse package (v1.3.0) with custom scripts available at the GitHub repository. Microchromosome and macrochromosome labels, when not available, were assigned to assembled sequences based on published karyotype data summarized in Table S1 and S2.

### GC content and gene density

GC content of each scaffold was calculated using BBMap (v38. 9) (65) for DNAzoo data, or obtained from the relevant NCBI genome information website. Gene density per megabase was calculated by dividing the number of annotated genes on a chromosome by its length.

### Interscaffold interactions

The HiC data were obtained from DNAzoo (Table S1). HiC matrices were exported to 50kb GInteractions bins with HiCExplorer (v3.6) (66). This format consisted of 7 columns: origin scaffold, origin scaffold start, origin scaffold end, target scaffold, target scaffold start, target scaffold end, and number of interactions between bins. The largest scaffolds were extracted, equal to the expected number of chromosomes based on the karyotype information (Table S1). Each square of the matrix represents the mean of normalized interaction values between the extracted scaffolds at 50kb bin resolution.

### Distance dependant contact probability P(s)

The HiC matrices were exported with HiCExplorer (v3.6) to h5 format. The scaffolds for each species were classified as either macro or microchromosomes (Table S2). Using the hicAdjustMatrix function, independent matrices were created for macrochromosomes and microchromosomes. Distance dependent contact probabilities [P(s)] were calculated using hicPlotDistVsCounts (from the HiCExplorer package), which were plotted with a maximum distance of 1×10^8^ bp.

### Golden eagle analysis

Regions of the golden eagle genome were classified according to homology with the chicken genome. These were: macro (macro in both species), micro (micro in both species), new macro (micro in chicken but macro in eagle) or new micro (macro in chicken but micro in eagle). These newly created scaffolds were then used to calculate GC content and distance dependent contact probabilities as described above.

## Acknowledgements

PK gratefully acknowledge the resources provided by University of Western Australia, Aiden lab at Baylor College of Medicine, DNA Zoo consortium partners and additional computational resources and support from the Pawsey Supercomputing Centre with funding from the Australian Government and the Government of Western Australia.

## References

1. J. M. van Brink, L’expression morphologique de la digametie chez les sauropsides et les monotremes. Chromosoma 10, 1–72 (1959).

2. L. Christidis, Animal Cytogenetics 4: Chordata 3. B. Aves (Gebrüder Borntraeger, Berlin, Germany).

3. N. Takagi, M. Sasaki, A phylogenetic study of bird karyotypes. Chromosoma 46, 91–120 (1974).

4. F. A. Habermann et al., Arrangements of macro-and microchromosomes in chicken cells. Chromosome Res 9, 569–584 (2001).

5. E. H. Newcomer, Accessory chromosomes in the domestic fowl. Genetics 40, 587–588 (1955).

6. H. A. McQueen, G. Siriaco, A. P. Bird, Chicken microchromosomes are hyperacetylated, early replicating, and gene rich. Genome Res 8, 621–630 (1998).

7. J. Smith et al., Differences in gene density on chicken macrochromosomes and microchromosomes. Anim Genet 31, 96–103 (2000).

8. D. W. Burt, Origin and evolution of avian microchromosomes. Cytogenet Genome Res 96, 97–112 (2002).

9. I. Nanda, M. Schmid, Localization of the telomeric (TTAGGG)n sequence in chicken (Gallus domesticus) chromosomes. Cytogenet Cell Genet 65, 190–193 (1994).

10. C. International Chicken Genome Sequencing, Sequence and comparative analysis of the chicken genome provide unique perspectives on vertebrate evolution. Nature 432, 695–716 (2004).

11. T. M. Degrandi et al., Introducing the Bird Chromosome Database: An Overview of Cytogenetic Studies in Birds. Cytogenet Genome Res 160, 199–205 (2020).

12. S. Shetty, D. K. Griffin, J. A. Graves, Comparative painting reveals strong chromosome homology over 80 million years of bird evolution. Chromosome Res 7, 289–295 (1999).

13. D. K. Griffin, L. B. Robertson, H. G. Tempest, B. M. Skinner, The evolution of the avian genome as revealed by comparative molecular cytogenetics. Cytogenet Genome Res 117, 64–77 (2007).

14. R. Kretschmer, M. A. Ferguson-Smith, E. H. C. de Oliveira, Karyotype Evolution in Birds: From Conventional Staining to Chromosome Painting. Genes (Basel) 9 (2018).

15. R. E. O’Connor et al., Patterns of microchromosome organization remain highly conserved throughout avian evolution. Chromosoma 128, 21–29 (2019).

16. R. Kretschmer et al., Interspecies Chromosome Mapping in Caprimulgiformes, Piciformes, Suliformes, and Trogoniformes (Aves): Cytogenomic Insight into Microchromosome Organization and Karyotype Evolution in Birds. Cells 10 (2021).

17. J. Damas, J. Kim, M. Farre, D. K. Griffin, D. M. Larkin, Reconstruction of avian ancestral karyotypes reveals differences in the evolutionary history of macro-and microchromosomes. Genome Biol 19, 155 (2018).

18. E. Olmo, Trends in the evolution of reptilian chromosomes. Integr Comp Biol 48, 486–493 (2008).

19. J. E. Deakin, T. Ezaz, Understanding the Evolution of Reptile Chromosomes through Applications of Combined Cytogenetics and Genomics Approaches. Cytogenet Genome Res 157, 7–20 (2019).

20. S. M. I. Alam, S. D. Sarre, D. Gleeson, A. Georges, T. Ezaz, Did Lizards Follow Unique Pathways in Sex Chromosome Evolution? Genes (Basel) 9 (2018).

21. Y. Matsuda et al., Highly conserved linkage homology between birds and turtles: bird and turtle chromosomes are precise counterparts of each other. Chromosome Res 13, 601–615 (2005).

22. W. Becak, M. L. Becak, H. R. Nazareth, S. Ohno, Close Karyological Kinship between the Reptilian Suborder Serpentes and the Class Aves. Chromosoma 15, 606–617 (1964).

23. Y. Uno et al., Inference of the protokaryotypes of amniotes and tetrapods and the evolutionary processes of microchromosomes from comparative gene mapping. PLoS One 7, e53027 (2012).

24. Z. Huang et al., Recurrent chromosome reshuffling and the evolution of neo-sex chromosomes in parrots. bioRxiv 10.1101/2021.03.08.434498, 2021.2003.2008.434498 (2021).

25. V. C. S. Oliveira et al., Revisiting the Karyotypes of Alligators and Caimans (Crocodylia, Alligatoridae) after a Half-Century Delay: Bridging the Gap in the Chromosomal Evolution of Reptiles. Cells 10, 1397 (2021).

26. S. R. Voss et al., Origin of amphibian and avian chromosomes by fission, fusion, and retention of ancestral chromosomes. Genome Res 21, 1306–1312 (2011).

27. A. Kapusta, A. Suh, Evolution of bird genomes-a transposon’s-eye view. Ann N Y Acad Sci 1389, 164–185 (2017).

28. Y. Nakatani, H. Takeda, Y. Kohara, S. Morishita, Reconstruction of the vertebrate ancestral genome reveals dynamic genome reorganization in early vertebrates. Genome Res 17, 1254–1265 (2007).

29. O. Simakov et al., Deeply conserved synteny resolves early events in vertebrate evolution. Nat Ecol Evol 4, 820–830 (2020).

30. G. Zhang et al., Comparative genomics reveals insights into avian genome evolution and adaptation. Science 346, 1311–1320 (2014).

31. R. S. Harris (2007) Improved pairwise alignment of genomic DNA. (The Pennsylvania State University).

32. M. Kohn, H. Kehrer-Sawatzki, W. Vogel, J. A. M. Graves, H. Hameister, Wide genome comparisons reveal the origins of the human X chromosome. Trends Genet 20, 598–603 (2004).

33. J. Liu et al., A new emu genome illuminates the evolution of genome configuration and nuclear architecture of avian chromosomes. Genome Res 31, 497–511 (2021).

34. R. A. Pyron, F. T. Burbrink, J. J. Wiens, A phylogeny and revised classification of Squamata, including 4161 species of lizards and snakes. BMC Evol Biol 13, 93 (2013).

35. A. Meyer et al., Giant lungfish genome elucidates the conquest of land by vertebrates. Nature 590, 284–289 (2021).

36. F. Kasai, P. C. O’Brien, M. A. Ferguson-Smith, Reassessment of genome size in turtle and crocodile based on chromosome measurement by flow karyotyping: close similarity to chicken. Biol Lett 8, 631–635 (2012).

37. D. R. Schield et al., The origins and evolution of chromosomes, dosage compensation, and mechanisms underlying venom regulation in snakes. Genome Res 29, 590–601 (2019).

38. B. W. Perry, D. R. Schield, R. H. Adams, T. A. Castoe, Microchromosomes Exhibit Distinct Features of Vertebrate Chromosome Structure and Function with Underappreciated Ramifications for Genome Evolution. Mol Biol Evol 38, 904–910 (2021).

39. O. Dudchenko et al., De novo assembly of the Aedes aegypti genome using Hi-C yields chromosome-length scaffolds. Science 356, 92–95 (2017).

40. B. R. Lajoie, J. Dekker, N. Kaplan, The Hitchhiker’s guide to Hi-C analysis: practical guidelines. Methods 72, 65–75 (2015).

41. J. Dekker, M. A. Marti-Renom, L. A. Mirny, Exploring the three-dimensional organization of genomes: interpreting chromatin interaction data. Nat Rev Genet 14, 390–403 (2013).

42. T. Raudsepp et al., Cytogenetic analysis of California condor (Gymnogyps californianus) chromosomes: comparison with chicken (Gallus gallus) macrochromosomes. Cytogenet Genome Res 98, 54–60 (2002).

43. D. McMillan et al., Characterizing the chromosomes of the platypus (Ornithorhynchus anatinus). Chromosome Res 15, 961–974 (2007).

44. T. D. Lamb, Analysis of Paralogons, Origin of the Vertebrate Karyotype, and Ancient Chromosomes Retained in Extant Species. Genome Biol Evol 13 (2021).

45. A. Canapa, M. Barucca, M. A. Biscotti, M. Forconi, E. Olmo, Transposons, Genome Size, and Evolutionary Insights in Animals. Cytogenet Genome Res 147, 217–239 (2015).

46. I. Nanda et al., Distribution of telomeric (TTAGGG)(n) sequences in avian chromosomes. Chromosoma 111, 215–227 (2002).

47. A. Burga et al., A genetic signature of the evolution of loss of flight in the Galapagos cormorant. Science 356 (2017).

48. J. Zhang, C. Yu, L. Krishnaswamy, T. Peterson, Transposable elements as catalysts for chromosome rearrangements. Methods Mol Biol 701, 315–326 (2011).

49. T. Marques-Bonet, A. Navarro, Chromosomal rearrangements are associated with higher rates of molecular evolution in mammals. Gene 353, 147–154 (2005).

50. Y. Zhang et al., Spatial organization of the mouse genome and its role in recurrent chromosomal translocations. Cell 148, 908–921 (2012).

51. D. Kordis, Transposable elements in reptilian and avian (sauropsida) genomes. Cytogenet Genome Res 127, 94–111 (2009).

52. T. Sultana, A. Zamborlini, G. Cristofari, P. Lesage, Integration site selection by retroviruses and transposable elements in eukaryotes. Nat Rev Genet 18, 292–308 (2017).

53. F. Grutzner et al., Chicken microchromosomes are hypermethylated and can be identified by specific painting probes. Cytogenet Cell Genet 93, 265–269 (2001).

54. G. Hartley, R. J. O’Neill, Centromere Repeats: Hidden Gems of the Genome. Genes (Basel) 10 (2019).

55. R. E. O’Connor et al., Chromosome-level assembly reveals extensive rearrangement in saker falcon and budgerigar, but not ostrich, genomes. Genome Biol 19, 171 (2018).

56. A. S. Graphodatsky, P. L. Perelman, S. J. O’Brien (2020) Atlas of mammalian chromosomes.

57. J. Kim et al., Reconstruction and evolutionary history of eutherian chromosomes. Proc Natl Acad Sci U S A 114, E5379–E5388 (2017).

58. M. Svartman, G. Stone, R. Stanyon, The ancestral eutherian karyotype is present in Xenarthra. PLoS Genet 2, e109 (2006).

59. J. E. Deakin et al., Reconstruction of the ancestral marsupial karyotype from comparative gene maps. BMC Evol Biol 13, 258 (2013).

60. S. J. Klein, R. J. O’Neill, Transposable elements: genome innovation, chromosome diversity, and centromere conflict. Chromosome Res 26, 5–23 (2018).

61. M. J. D. White, Modes of Speciation (W. H. Freeman, San Francisco, 1978).

62. T. Cremer, C. Cremer, Chromosome territories, nuclear architecture and gene regulation in mammalian cells. Nat Rev Genet 2, 292–301 (2001).

63. T. Ezaz et al., The dragon lizard Pogona vitticeps has ZZ/ZW micro-sex chromosomes. Chromosome Res 13, 763–776 (2005).

64. K. Matsubara et al., Molecular cloning and characterization of satellite DNA sequences from constitutive heterochromatin of the habu snake (Protobothrops flavoviridis, Viperidae) and the Burmese python (Python bivittatus, Pythonidae). Chromosoma 124, 529–539 (2015).

65. B. Bushnell (2014) BBMap: A Fast, Accurate, Splice-Aware Aligner.

66. J. Wolff et al., Galaxy HiCExplorer 3: a web server for reproducible Hi-C, capture Hi-C and single-cell Hi-C data analysis, quality control and visualization. Nucleic Acids Res 48, W177–W184 (2020).

