## Supplementary Information for "Microchromosomes are building blocks of bird, reptile and mammal chromosomes"

Jennifer A. Marshall Graves

**This PDF file includes:**

Figures S1 to S3

Tables S1 to S2

SI References

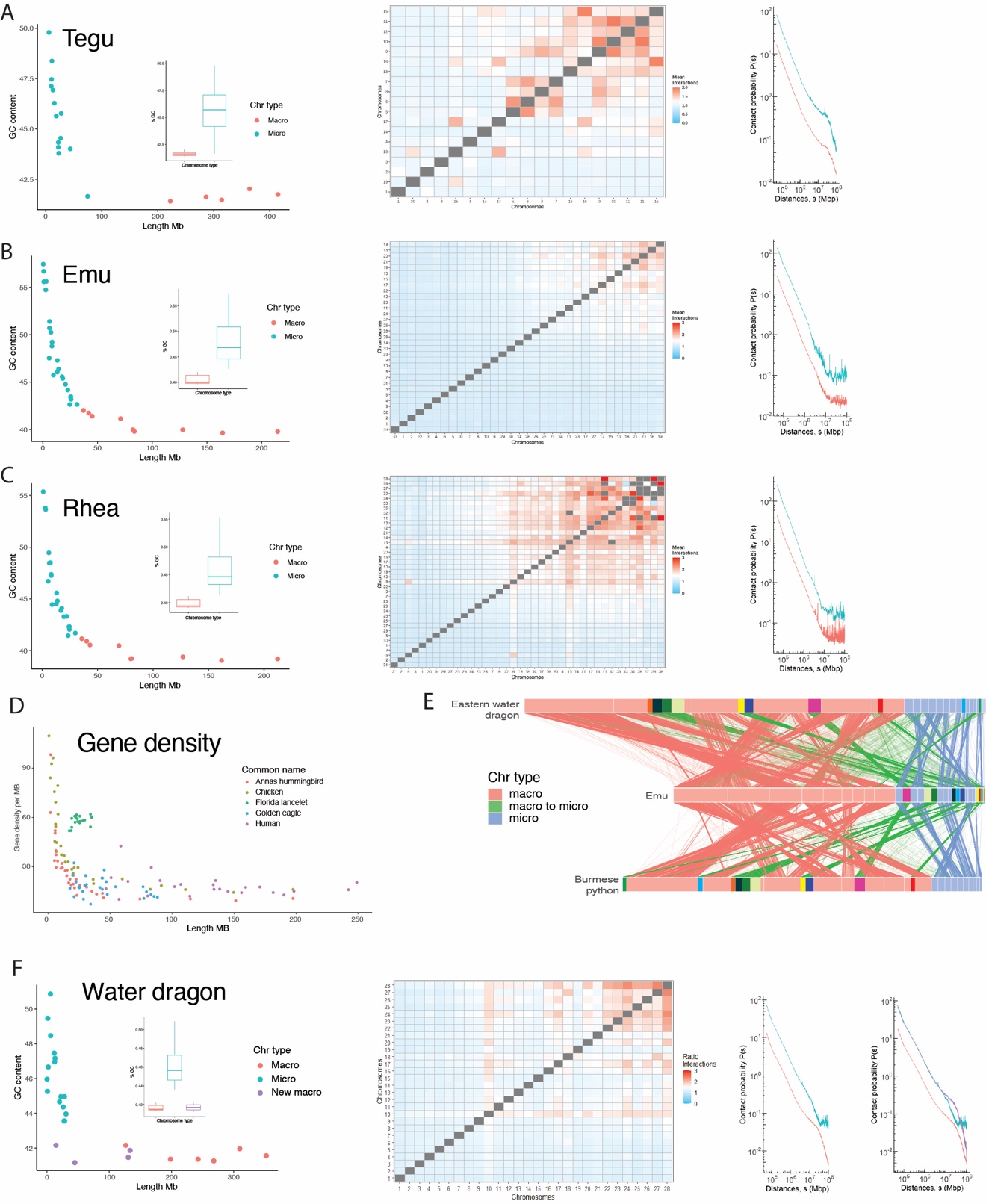

Fig. S1. GC content versus chromosome size, inter-chromosome interaction ratios (with scaffolds sorted from largest to smallest) and distance dependent interaction probabilities of macro and microchromosomes (Ps) for: (A) tegu, (B) emu, (C) rhea. (D) Gene density versus chromosome size for representative bird species along with Florida lancelet and human. (E) Incorporation of bird (emu) microchromosomes into the water dragon and python macrochromosome. Each emu microchromosome incorporated into a macrochromosome in colored differently. (F) GC content versus chromosome size, inter-chromosome interaction ratios (with scaffolds sorted from largest to smallest) and distance dependent interaction probabilities of macro and microchromosomes (Ps) for water dragon. Water dragon has four regions of macrochromosomes that are homologous to ancestral microchromosomes (see panel E). These new macrochromosome regions have reduced GC content, but maintain high contact probabilities.

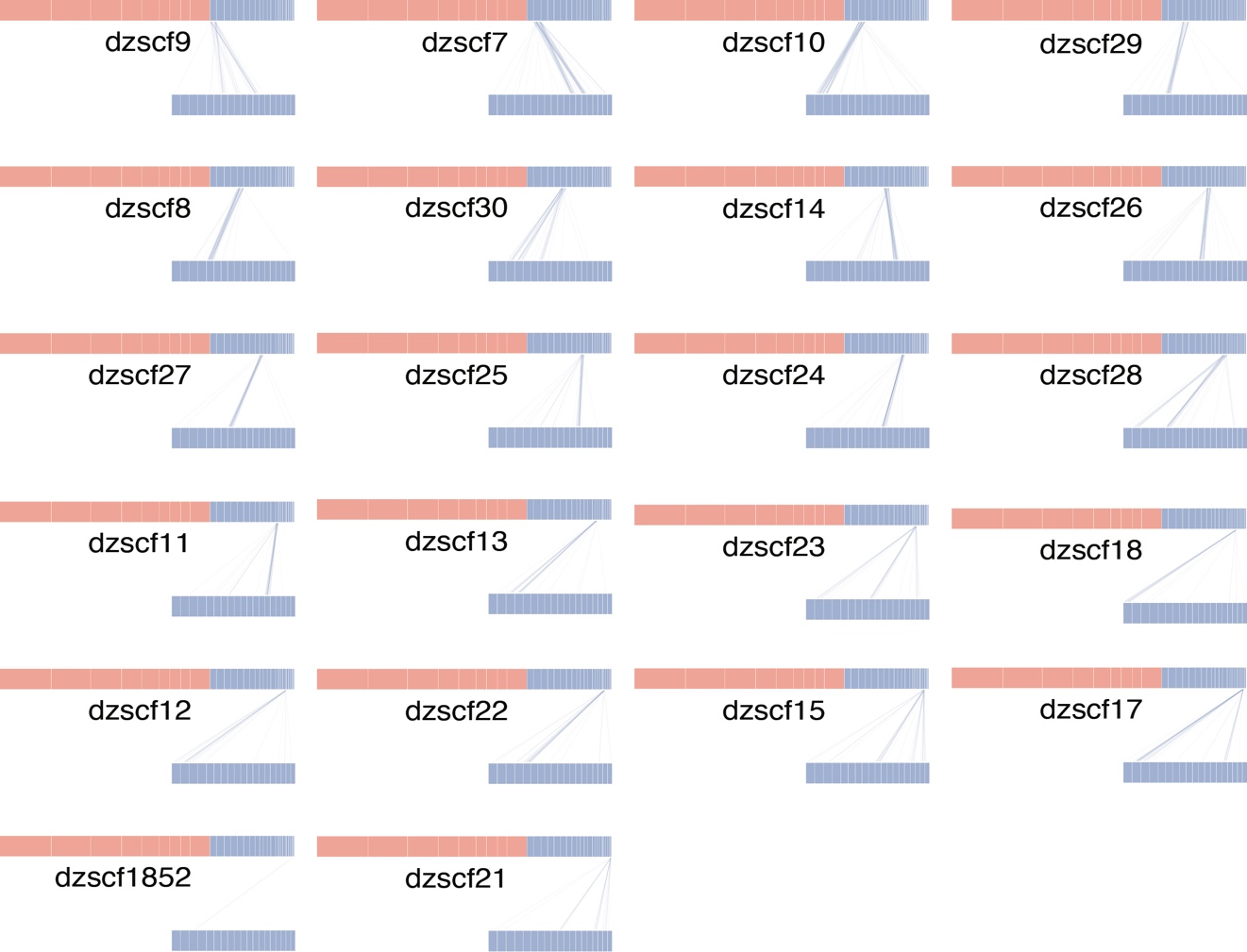

Fig. S2. Homologies of individual emu chromosomes to the amphioxus genome. Chained and netted alignments were filtered for a minimum length of 5kb.

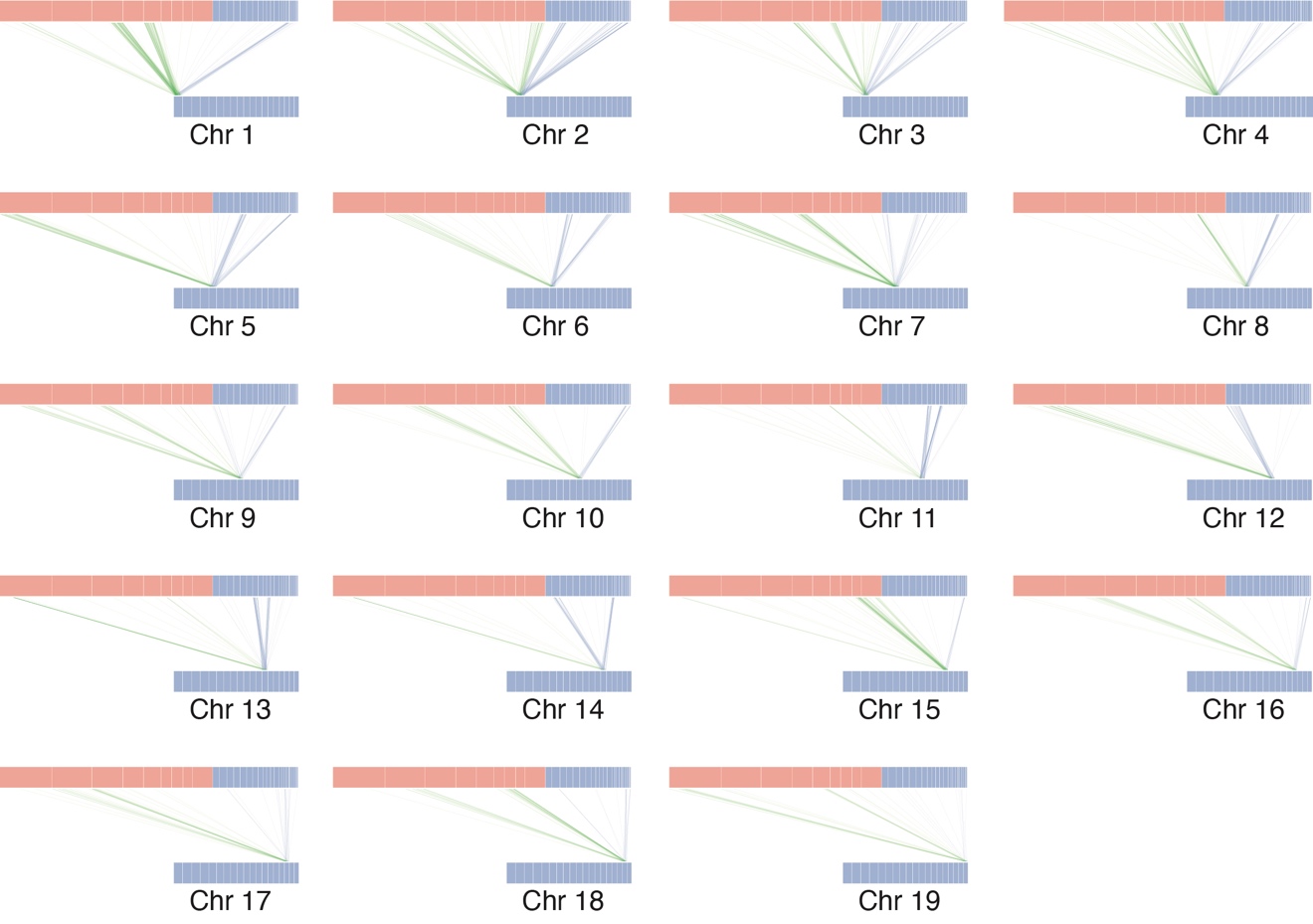

Fig. S3. Homologies of individual amphioxus chromosomes to the emu genome. Homologies to macrochromosomes are green. Homologies to microchromosomes are blue. Chained and netted alignments were filtered for a minimum length of 5kb.

Table S1. Information about species included in this study: Scientific name, common name, links to sequence data, references for karyotypes information, cytological chromosome number, number of assembled macrochromosomes, number of assembled microchromosomes, smallest assembled macrochromosome, and largest assembled microchromosome.

| **Species** | **Common name** | **Sequence fasta file address** | **Karyotype reference** | **Chr number (n)** | **Macro assembled (n)** | **Micro assembled (n)** | **Smallest macro** | **Largest micro** | **Notes** |
| --- | --- | --- | --- | --- | --- | --- | --- | --- | --- |
| Podarcis muralis | Common wall lizard | https://ftp.ncbi.nlm.nih.gov/genomes/all/GCF/004/329/235/GCF_004329235.1_PodMur_1.0/GCF_004329235.1_PodMur_1.0_genomic.fna.gz | (1) | 19 | 15 | 4 | 54988359 | 46441056 | No clear macro micro distinction in the karyotype. 50Mb threshold chosen. |
| Zootoca vivipara | Common lizard | https://ftp.ncbi.nlm.nih.gov/genomes/all/GCF/011/800/845/GCF_011800845.1_UG_Zviv_1/GCF_011800845.1_UG_Zviv_1_genomic.fna.gz | (2) | 19 | 11 | 8 | 51059636 | 49589567 | No clear macro micro distinction in the karyotype. 50Mb threshold chosen. |
| Salvator merianae | Argentine black and white tegu | https://www.dropbox.com/s/x6vut6u55osw7pt/HLtupMer6_HiC.fasta.gz | (3) | 19 | 5 | 14 | 222499501 | 74934432 |  |
| Intellagama lesueurii lesueurii | Eastern water dragon | <https://www.dropbox.com/s/kwhifyc2ok7r7mw/EWD_hifiasm_HiC.fasta.gz> | NA | ? | 6 | 19 | 127569639 | 31587450 | No karyotype. |
| Python bivittatus | Burmese python | https://www.dropbox.com/s/rzucyl0nkjnt12a/Python_molurus_bivittatus-5.0.2_HiC.fasta.gz | (4) | 18 | 8 | 10 | 72243982 | 25188946 |  |
| Crotalus viridis | Prairie rattlesnake | https://ftp.ncbi.nlm.nih.gov/genomes/all/GCA/003/400/415/GCA_003400415.2_UTA_CroVir_3.0/GCA_003400415.2_UTA_CroVir_3.0_genomic.fna.gz | (5) | 19 | 8 | 11 | 61662597 | 22521304 |  |
| Naja naja | Indian cobra | https://ftp.ncbi.nlm.nih.gov/genomes/all/GCA/009/733/165/GCA_009733165.1_Nana_v5/GCA_009733165.1_Nana_v5_genomic.fna.gz | (6) | 19 | 8 | 11 | 53796468 | 32026552 |  |
| Dromaius novaehollandiae | Emu | https://www.dropbox.com/s/bge457t6l9a5uqp/droNov1_HiC.fasta.gz | (7) | 40 | 9 | 31 | 37106476 | 31258799 |  |
| Rhea americana | Greater rhea | https://www.dropbox.com/s/c52npppt0m7e2dv/rheAme1_HiC.fasta.gz | (8) | 40 | 9 | 21 | 35777221 | 29910934 |  |
| Gallus gallus | Chicken | https://ftp.ncbi.nlm.nih.gov/genomes/all/GCF/000/002/315/GCF_000002315.6_GRCg6a/GCF_000002315.6_GRCg6a_genomic.fna.gz | (9) | 39 | 9 | 24 | 30219446 | 24153086 |  |
| Calypte anna | Anna's hummingbird | https://ftp.ncbi.nlm.nih.gov/genomes/all/GCF/003/957/555/GCF_003957555.1_bCalAnn1_v1.p/GCF_003957555.1_bCalAnn1_v1.p_genomic.fna.gz | (10) | 37 | 9 | 26 | 31090148 | 25686456 |  |
| Corvus monedula | Jackdaw | https://ftp.ncbi.nlm.nih.gov/genomes/all/GCA/013/407/035/GCA_013407035.1_ASM1340703v1/GCA_013407035.1_ASM1340703v1_genomic.fna.gz | (11) | 40 | 10 | 21 | 32060782 | 26026359 | No karyotype. Common crow has 2n = 80 |
| Patagioenas fasciata | Band-tailed pigeon | https://www.dropbox.com/s/kfmoz7kwtcuq44m/NIATT_ARIZONA_HiC.fasta.gz | (12) | 40 | 9 | 21 | 32469999 | 26560337 | No karyotype. Other pigeons 2n = 80 |
| Phalacrocorax auritus | Double-crested cormorant | https://www.dropbox.com/s/fh23wbsipgzniml/ASM217345v1_HiC.fasta.gz | (13) | 36-43 | 10 | 23 | 46711434 | 26678942 | No karyotype. Other species from the genus Phalacrocorax have 2n = 72-86 |
| Aquila chrysaetos chrysaetos | Golden eagle | https://www.dropbox.com/s/ynqlwde2k8xcijl/Aquila_chrysaetos-1.0.2_HiC.fasta.gz | (14) | 31 | 13 | 14 | 43481500 | 34337559 | No clear macro micro distinction in the karyotype. 40Mb threshold chosen. |
| Alligator mississippiensis | American alligator | https://www.dropbox.com/s/xdoke3xiyzxbkfb/ASM28112v4_HiC.fasta.gz | (15) | 16 | 5 | 10 | 209376224 | 95926144 |  |
| Chelonia mydas | Green sea turtle | <https://www.dropbox.com/s/x4zbv0mosgda9k8/CheMyd_1.0_HiC.fasta.gz> | (16) | 28 | 11 | 17 | 78168177 | 42804224 |  |
| Trachemys scripta elegans | Red-eared slider turtle | https://ftp.ncbi.nlm.nih.gov/genomes/all/GCF/013/100/865/GCF_013100865.1_CAS_Tse_1.0/GCF_013100865.1_CAS_Tse_1.0_genomic.fna.gz | (17) | 25 | 11 | 14 | 76047897 | 45061502 |  |
| Ornithorhynchus anatinus | Platypus | https://ftp.ncbi.nlm.nih.gov/genomes/refseq/vertebrate_mammalian/Ornithorhynchus_anatinus/latest_assembly_versions/GCF_004115215.1_mOrnAna1.p.v1/GCF_004115215.1_mOrnAna1.p.v1_genomic.fna.gz | (18) | 26 | NA | NA | NA | NA |  |
| Homo sapiens | Human | https://ftp.ncbi.nlm.nih.gov/genomes/all/GCA/000/001/405/GCA_000001405.28_GRCh38.p13/GRCh38_major_release_seqs_for_alignment_pipelines/GCA_000001405.15_GRCh38_no_alt_analysis_set.fna.gz | (19) | 23 | NA | NA | NA | NA |  |
| Phascolarctos cinereus | Koala | https://www.dropbox.com/s/66hp8hjonljvm3q/phaCin_unsw_v4.1_HiC.fasta.gz | (20) | 8 | NA | NA | NA | NA |  |

Table S2. Chromosome information and plot order for all species.

| Sequence ID | Scaffold length | Chromosome type | Position in plot | Species code | Chromosome name | Common name |
| --- | --- | --- | --- | --- | --- | --- |
| seqid | seqlength | chrtype | krank | speciescode | chrname | cname |
| ALLMI_HiC_scaffold_1 | 446151378 | macro | 1 | ALLMI | dzscf1 | American alligator |
| ALLMI_HiC_scaffold_2 | 291816272 | macro | 2 | ALLMI | dzscf2 | American alligator |
| ALLMI_HiC_scaffold_16 | 286864427 | macro | 3 | ALLMI | dzscf16 | American alligator |
| ALLMI_HiC_scaffold_3 | 240791973 | macro | 4 | ALLMI | dzscf3 | American alligator |
| ALLMI_HiC_scaffold_15 | 209376224 | macro | 5 | ALLMI | dzscf15 | American alligator |
| ALLMI_HiC_scaffold_14 | 95926144 | micro | 6 | ALLMI | dzscf14 | American alligator |
| ALLMI_HiC_scaffold_6 | 79122728 | micro | 8 | ALLMI | dzscf6 | American alligator |
| ALLMI_HiC_scaffold_4 | 77868838 | micro | 7 | ALLMI | dzscf4 | American alligator |
| ALLMI_HiC_scaffold_7 | 73747066 | micro | 9 | ALLMI | dzscf7 | American alligator |
| ALLMI_HiC_scaffold_5 | 71713467 | micro | 10 | ALLMI | dzscf5 | American alligator |
| ALLMI_HiC_scaffold_12 | 65528272 | micro | 11 | ALLMI | dzscf12 | American alligator |
| ALLMI_HiC_scaffold_8 | 56766657 | micro | 13 | ALLMI | dzscf8 | American alligator |
| ALLMI_HiC_scaffold_13 | 54226289 | micro | 12 | ALLMI | dzscf13 | American alligator |
| ALLMI_HiC_scaffold_11 | 36339844 | micro | 14 | ALLMI | dzscf11 | American alligator |
| ALLMI_HiC_scaffold_9 | 27889950 | micro | 16 | ALLMI | dzscf9 | American alligator |
| ALLMI_HiC_scaffold_10 | 23283895 | micro | 15 | ALLMI | dzscf10 | American alligator |
| DRONO_HiC_scaffold_33 | 214659357 | macro | 1 | DRONO | dzscf33 | Emu |
| DRONO_HiC_scaffold_1 | 164242004 | macro | 2 | DRONO | dzscf1 | Emu |
| DRONO_HiC_scaffold_2 | 127852514 | macro | 3 | DRONO | dzscf2 | Emu |
| DRONO_HiC_scaffold_32 | 83679895 | macro | 4 | DRONO | dzscf32 | Emu |
| DRONO_HiC_scaffold_3 | 82460724 | macro | 9 | DRONO | dzscf3 | Emu |
| DRONO_HiC_scaffold_4 | 71046951 | macro | 5 | DRONO | dzscf4 | Emu |
| DRONO_HiC_scaffold_6 | 45086358 | macro | 7 | DRONO | dzscf6 | Emu |
| DRONO_HiC_scaffold_5 | 41922228 | macro | 6 | DRONO | dzscf5 | Emu |
| DRONO_HiC_scaffold_31 | 37106476 | macro | 8 | DRONO | dzscf31 | Emu |
| DRONO_HiC_scaffold_7 | 31258799 | micro | 11 | DRONO | dzscf7 | Emu |
| DRONO_HiC_scaffold_8 | 25801502 | micro | 14 | DRONO | dzscf8 | Emu |
| DRONO_HiC_scaffold_10 | 25601094 | micro | 12 | DRONO | dzscf10 | Emu |
| DRONO_HiC_scaffold_9 | 25287843 | micro | 10 | DRONO | dzscf9 | Emu |
| DRONO_HiC_scaffold_29 | 24679445 | micro | 13 | DRONO | dzscf29 | Emu |
| DRONO_HiC_scaffold_30 | 22865478 | micro | 15 | DRONO | dzscf30 | Emu |
| DRONO_HiC_scaffold_14 | 20727510 | micro | 16 | DRONO | dzscf14 | Emu |
| DRONO_HiC_scaffold_28 | 18241439 | micro | 21 | DRONO | dzscf28 | Emu |
| DRONO_HiC_scaffold_26 | 17562204 | micro | 17 | DRONO | dzscf26 | Emu |
| DRONO_HiC_scaffold_25 | 14364803 | micro | 19 | DRONO | dzscf25 | Emu |
| DRONO_HiC_scaffold_27 | 13296474 | micro | 18 | DRONO | dzscf27 | Emu |
| DRONO_HiC_scaffold_24 | 13249401 | micro | 20 | DRONO | dzscf24 | Emu |
| DRONO_HiC_scaffold_11 | 9663520 | micro | 22 | DRONO | dzscf11 | Emu |
| DRONO_HiC_scaffold_23 | 8479663 | micro | 24 | DRONO | dzscf23 | Emu |
| DRONO_HiC_scaffold_12 | 8297778 | micro | 25 | DRONO | dzscf12 | Emu |
| DRONO_HiC_scaffold_22 | 7430024 | micro | 27 | DRONO | dzscf22 | Emu |
| DRONO_HiC_scaffold_17 | 6134115 | micro | 29 | DRONO | dzscf17 | Emu |
| DRONO_HiC_scaffold_15 | 6066571 | micro | 28 | DRONO | dzscf15 | Emu |
| DRONO_HiC_scaffold_13 | 5805633 | micro | 23 | DRONO | dzscf13 | Emu |
| DRONO_HiC_scaffold_18 | 3111746 | micro | 26 | DRONO | dzscf18 | Emu |
| DRONO_HiC_scaffold_21 | 2652061 | micro | 31 | DRONO | dzscf21 | Emu |
| DRONO_HiC_scaffold_1852 | 1933103 | micro | 30 | DRONO | dzscf1852 | Emu |
| PYTBI_HiC_scaffold_2 | 317910269 | macro | 1 | PYTBI | dzscf2 | Burmese python |
| PYTBI_HiC_scaffold_1 | 241438387 | macro | 2 | PYTBI | dzscf1 | Burmese python |
| PYTBI_HiC_scaffold_6 | 196651090 | macro | 3 | PYTBI | dzscf6 | Burmese python |
| PYTBI_HiC_scaffold_4 | 108932528 | macro | 4 | PYTBI | dzscf4 | Burmese python |
| PYTBI_HiC_scaffold_5 | 106496723 | macro | 8 | PYTBI | dzscf5 | Burmese python |
| PYTBI_HiC_scaffold_18 | 93335423 | macro | 5 | PYTBI | dzscf18 | Burmese python |
| PYTBI_HiC_scaffold_7 | 78792744 | macro | 7 | PYTBI | dzscf7 | Burmese python |
| PYTBI_HiC_scaffold_8 | 72243982 | macro | 6 | PYTBI | dzscf8 | Burmese python |
| PYTBI_HiC_scaffold_15 | 25188946 | micro | 10 | PYTBI | dzscf15 | Burmese python |
| PYTBI_HiC_scaffold_3 | 24620612 | micro | 12 | PYTBI | dzscf3 | Burmese python |
| PYTBI_HiC_scaffold_17 | 23955143 | micro | 9 | PYTBI | dzscf17 | Burmese python |
| PYTBI_HiC_scaffold_16 | 22221336 | micro | 11 | PYTBI | dzscf16 | Burmese python |
| PYTBI_HiC_scaffold_9 | 21648691 | micro | 13 | PYTBI | dzscf9 | Burmese python |
| PYTBI_HiC_scaffold_12 | 18726175 | micro | 14 | PYTBI | dzscf12 | Burmese python |
| PYTBI_HiC_scaffold_10 | 15207260 | micro | 15 | PYTBI | dzscf10 | Burmese python |
| PYTBI_HiC_scaffold_11 | 11460064 | micro | 16 | PYTBI | dzscf11 | Burmese python |
| PYTBI_HiC_scaffold_13 | 9923712 | micro | 17 | PYTBI | dzscf13 | Burmese python |
| PYTBI_HiC_scaffold_14 | 9400649 | micro | 18 | PYTBI | dzscf14 | Burmese python |
| SAMER_HiC_scaffold_1 | 414523068 | macro | 1 | SAMER | dzscf1 | Argentine black and white tegu |
| SAMER_HiC_scaffold_19 | 363964080 | macro | 2 | SAMER | dzscf19 | Argentine black and white tegu |
| SAMER_HiC_scaffold_18 | 314194259 | macro | 4 | SAMER | dzscf18 | Argentine black and white tegu |
| SAMER_HiC_scaffold_2 | 286286133 | macro | 3 | SAMER | dzscf2 | Argentine black and white tegu |
| SAMER_HiC_scaffold_3 | 222499501 | macro | 5 | SAMER | dzscf3 | Argentine black and white tegu |
| SAMER_HiC_scaffold_17 | 74934432 | micro | 7 | SAMER | dzscf17 | Argentine black and white tegu |
| SAMER_HiC_scaffold_16 | 43927702 | micro | 6 | SAMER | dzscf16 | Argentine black and white tegu |
| SAMER_HiC_scaffold_15 | 27733050 | micro | 8 | SAMER | dzscf15 | Argentine black and white tegu |
| SAMER_HiC_scaffold_5 | 26807181 | micro | 9 | SAMER | dzscf5 | Argentine black and white tegu |
| SAMER_HiC_scaffold_4 | 23377602 | micro | 10 | SAMER | dzscf4 | Argentine black and white tegu |
| SAMER_HiC_scaffold_6 | 22704898 | micro | 11 | SAMER | dzscf6 | Argentine black and white tegu |
| SAMER_HiC_scaffold_7 | 22666949 | micro | 12 | SAMER | dzscf7 | Argentine black and white tegu |
| SAMER_HiC_scaffold_9 | 18916873 | micro | 13 | SAMER | dzscf9 | Argentine black and white tegu |
| SAMER_HiC_scaffold_8 | 15552402 | micro | 14 | SAMER | dzscf8 | Argentine black and white tegu |
| SAMER_HiC_scaffold_10 | 12965002 | micro | 15 | SAMER | dzscf10 | Argentine black and white tegu |
| SAMER_HiC_scaffold_11 | 11229012 | micro | 16 | SAMER | dzscf11 | Argentine black and white tegu |
| SAMER_HiC_scaffold_13 | 10575517 | micro | 17 | SAMER | dzscf13 | Argentine black and white tegu |
| SAMER_HiC_scaffold_14 | 10004368 | micro | 18 | SAMER | dzscf14 | Argentine black and white tegu |
| SAMER_HiC_scaffold_12 | 5890402 | micro | 19 | SAMER | dzscf12 | Argentine black and white tegu |
| HiC_scaffold_1 | 730168553 | macro | 1 | PHACI | dzscf1 | Koala |
| HiC_scaffold_2 | 630767262 | macro | 2 | PHACI | dzscf2 | Koala |
| HiC_scaffold_3 | 480107551 | macro | 3 | PHACI | dzscf3 | Koala |
| HiC_scaffold_4 | 413949655 | macro | 4 | PHACI | dzscf4 | Koala |
| HiC_scaffold_5 | 294865928 | macro | 5 | PHACI | dzscf5 | Koala |
| HiC_scaffold_6 | 265033287 | macro | 6 | PHACI | dzscf6 | Koala |
| HiC_scaffold_7 | 260546985 | macro | 7 | PHACI | dzscf7 | Koala |
| HiC_scaffold_8 | 75917611 | macro | 8 | PHACI | dzscf8 | Koala |
| HiC_scaffold_1 | 86574484 | macro | 20 | AQUCHDZ | dzscf1 | Golden eagle DNAZOO |
| HiC_scaffold_2 | 85437313 | macro | 14 | AQUCHDZ | dzscf2 | Golden eagle DNAZOO |
| HiC_scaffold_3 | 82400852 | macro | 15 | AQUCHDZ | dzscf3 | Golden eagle DNAZOO |
| HiC_scaffold_4 | 79048268 | macro | 10 | AQUCHDZ | dzscf4 | Golden eagle DNAZOO |
| HiC_scaffold_5 | 77143528 | macro | 8 | AQUCHDZ | dzscf5 | Golden eagle DNAZOO |
| HiC_scaffold_6 | 75989677 | macro | 1 | AQUCHDZ | dzscf6 | Golden eagle DNAZOO |
| HiC_scaffold_7 | 54380017 | macro | 18 | AQUCHDZ | dzscf7 | Golden eagle DNAZOO |
| HiC_scaffold_8 | 47735502 | macro | 4 | AQUCHDZ | dzscf8 | Golden eagle DNAZOO |
| HiC_scaffold_9 | 46579469 | macro | 13 | AQUCHDZ | dzscf9 | Golden eagle DNAZOO |
| HiC_scaffold_10 | 44400008 | macro | 23 | AQUCHDZ | dzscf10 | Golden eagle DNAZOO |
| HiC_scaffold_11 | 43719360 | macro | 22 | AQUCHDZ | dzscf11 | Golden eagle DNAZOO |
| HiC_scaffold_12 | 43294077 | macro | 19 | AQUCHDZ | dzscf12 | Golden eagle DNAZOO |
| HiC_scaffold_13 | 43238996 | macro | 17 | AQUCHDZ | dzscf13 | Golden eagle DNAZOO |
| HiC_scaffold_14 | 42114285 | macro | 12 | AQUCHDZ | dzscf14 | Golden eagle DNAZOO |
| HiC_scaffold_15 | 34286477 | micro | 6 | AQUCHDZ | dzscf15 | Golden eagle DNAZOO |
| HiC_scaffold_16 | 30665523 | micro | 11 | AQUCHDZ | dzscf16 | Golden eagle DNAZOO |
| HiC_scaffold_17 | 30406229 | micro | 16 | AQUCHDZ | dzscf17 | Golden eagle DNAZOO |
| HiC_scaffold_18 | 29672543 | micro | 3 | AQUCHDZ | dzscf18 | Golden eagle DNAZOO |
| HiC_scaffold_19 | 28594004 | micro | 9 | AQUCHDZ | dzscf19 | Golden eagle DNAZOO |
| HiC_scaffold_20 | 27848399 | micro | 7 | AQUCHDZ | dzscf20 | Golden eagle DNAZOO |
| HiC_scaffold_21 | 24930757 | micro | 24 | AQUCHDZ | dzscf21 | Golden eagle DNAZOO |
| HiC_scaffold_22 | 24705485 | micro | 21 | AQUCHDZ | dzscf22 | Golden eagle DNAZOO |
| HiC_scaffold_23 | 22133620 | micro | 25 | AQUCHDZ | dzscf23 | Golden eagle DNAZOO |
| HiC_scaffold_24 | 20915179 | micro | 27 | AQUCHDZ | dzscf24 | Golden eagle DNAZOO |
| HiC_scaffold_25 | 20589069 | micro | 5 | AQUCHDZ | dzscf25 | Golden eagle DNAZOO |
| HiC_scaffold_26 | 19774316 | micro | 26 | AQUCHDZ | dzscf26 | Golden eagle DNAZOO |
| HiC_scaffold_27 | 17479260 | micro | 2 | AQUCHDZ | dzscf27 | Golden eagle DNAZOO |
| HiC_scaffold_28 | 1424811 | micro | 28 | AQUCHDZ | dzscf28 | Golden eagle DNAZOO |
| HiC_scaffold_1 | 344580086 | macro | 1 | CHEMYDZ | dzscf1 | Green sea turtle DNAZOO |
| HiC_scaffold_2 | 265960234 | macro | 2 | CHEMYDZ | dzscf2 | Green sea turtle DNAZOO |
| HiC_scaffold_17 | 206018961 | macro | 3 | CHEMYDZ | dzscf17 | Green sea turtle DNAZOO |
| HiC_scaffold_24 | 140825401 | macro | 4 | CHEMYDZ | dzscf24 | Green sea turtle DNAZOO |
| HiC_scaffold_18 | 133491730 | macro | 9 | CHEMYDZ | dzscf18 | Green sea turtle DNAZOO |
| HiC_scaffold_20 | 127535543 | macro | 5 | CHEMYDZ | dzscf20 | Green sea turtle DNAZOO |
| HiC_scaffold_3 | 125259731 | macro | 6 | CHEMYDZ | dzscf3 | Green sea turtle DNAZOO |
| HiC_scaffold_4 | 107487999 | macro | 8 | CHEMYDZ | dzscf4 | Green sea turtle DNAZOO |
| HiC_scaffold_25 | 102862947 | macro | 11 | CHEMYDZ | dzscf25 | Green sea turtle DNAZOO |
| HiC_scaffold_23 | 85189569 | macro | 10 | CHEMYDZ | dzscf23 | Green sea turtle DNAZOO |
| HiC_scaffold_19 | 78168177 | macro | 7 | CHEMYDZ | dzscf19 | Green sea turtle DNAZOO |
| HiC_scaffold_11 | 42804224 | micro | 12 | CHEMYDZ | dzscf11 | Green sea turtle DNAZOO |
| HiC_scaffold_10 | 39090782 | micro | 17 | CHEMYDZ | dzscf10 | Green sea turtle DNAZOO |
| HiC_scaffold_5 | 39009518 | micro | 15 | CHEMYDZ | dzscf5 | Green sea turtle DNAZOO |
| HiC_scaffold_22 | 33173454 | micro | 13 | CHEMYDZ | dzscf22 | Green sea turtle DNAZOO |
| HiC_scaffold_26 | 26159407 | micro | 14 | CHEMYDZ | dzscf26 | Green sea turtle DNAZOO |
| HiC_scaffold_7 | 25598500 | micro | 16 | CHEMYDZ | dzscf7 | Green sea turtle DNAZOO |
| HiC_scaffold_15 | 23668442 | micro | 18 | CHEMYDZ | dzscf15 | Green sea turtle DNAZOO |
| HiC_scaffold_14 | 19940459 | micro | 20 | CHEMYDZ | dzscf14 | Green sea turtle DNAZOO |
| HiC_scaffold_12 | 19221767 | micro | 21 | CHEMYDZ | dzscf12 | Green sea turtle DNAZOO |
| HiC_scaffold_16 | 18802053 | micro | 23 | CHEMYDZ | dzscf16 | Green sea turtle DNAZOO |
| HiC_scaffold_21 | 17385582 | micro | 27 | CHEMYDZ | dzscf21 | Green sea turtle DNAZOO |
| HiC_scaffold_6 | 17094346 | micro | 22 | CHEMYDZ | dzscf6 | Green sea turtle DNAZOO |
| HiC_scaffold_13 | 16651426 | micro | 25 | CHEMYDZ | dzscf13 | Green sea turtle DNAZOO |
| HiC_scaffold_9 | 16544897 | micro | 19 | CHEMYDZ | dzscf9 | Green sea turtle DNAZOO |
| HiC_scaffold_8 | 16464228 | micro | 24 | CHEMYDZ | dzscf8 | Green sea turtle DNAZOO |
| HiC_scaffold_28 | 15684472 | micro | 26 | CHEMYDZ | dzscf28 | Green sea turtle DNAZOO |
| HiC_scaffold_27 | 5477977 | micro | 28 | CHEMYDZ | dzscf27 | Green sea turtle DNAZOO |
| HiC_scaffold_1 | 205510306 | macro | 1 | PATFA | dzscf1 | Band-tailed pigeon |
| HiC_scaffold_2 | 157166770 | macro | 2 | PATFA | dzscf2 | Band-tailed pigeon |
| HiC_scaffold_3 | 117214671 | macro | 3 | PATFA | dzscf3 | Band-tailed pigeon |
| HiC_scaffold_4 | 76042635 | macro | 9 | PATFA | dzscf4 | Band-tailed pigeon |
| HiC_scaffold_5 | 75366177 | macro | 4 | PATFA | dzscf5 | Band-tailed pigeon |
| HiC_scaffold_6 | 66896167 | macro | 5 | PATFA | dzscf6 | Band-tailed pigeon |
| HiC_scaffold_7 | 39534137 | macro | 7 | PATFA | dzscf7 | Band-tailed pigeon |
| HiC_scaffold_8 | 37860466 | macro | 6 | PATFA | dzscf8 | Band-tailed pigeon |
| HiC_scaffold_9 | 32469999 | macro | 8 | PATFA | dzscf9 | Band-tailed pigeon |
| HiC_scaffold_10 | 26560337 | micro | 11 | PATFA | dzscf10 | Band-tailed pigeon |
| HiC_scaffold_11 | 21829948 | micro | 14 | PATFA | dzscf11 | Band-tailed pigeon |
| HiC_scaffold_12 | 21065616 | micro | 12 | PATFA | dzscf12 | Band-tailed pigeon |
| HiC_scaffold_13 | 20908936 | micro | 10 | PATFA | dzscf13 | Band-tailed pigeon |
| HiC_scaffold_14 | 20716055 | micro | 13 | PATFA | dzscf14 | Band-tailed pigeon |
| HiC_scaffold_15 | 19177021 | micro | 15 | PATFA | dzscf15 | Band-tailed pigeon |
| HiC_scaffold_16 | 16763932 | micro | 16 | PATFA | dzscf16 | Band-tailed pigeon |
| HiC_scaffold_17 | 14726480 | micro | 21 | PATFA | dzscf17 | Band-tailed pigeon |
| HiC_scaffold_18 | 13961864 | micro | 17 | PATFA | dzscf18 | Band-tailed pigeon |
| HiC_scaffold_19 | 11361449 | micro | 19 | PATFA | dzscf19 | Band-tailed pigeon |
| HiC_scaffold_20 | 10997106 | micro | 18 | PATFA | dzscf20 | Band-tailed pigeon |
| HiC_scaffold_21 | 10252728 | micro | 20 | PATFA | dzscf21 | Band-tailed pigeon |
| HiC_scaffold_22 | 7012735 | micro | 22 | PATFA | dzscf22 | Band-tailed pigeon |
| HiC_scaffold_23 | 5841358 | micro | 26 | PATFA | dzscf23 | Band-tailed pigeon |
| HiC_scaffold_24 | 5675082 | micro | 25 | PATFA | dzscf24 | Band-tailed pigeon |
| HiC_scaffold_25 | 5607113 | micro | 24 | PATFA | dzscf25 | Band-tailed pigeon |
| HiC_scaffold_26 | 5091844 | micro | 27 | PATFA | dzscf26 | Band-tailed pigeon |
| HiC_scaffold_27 | 4504171 | micro | 28 | PATFA | dzscf27 | Band-tailed pigeon |
| HiC_scaffold_28 | 4433014 | micro | 23 | PATFA | dzscf28 | Band-tailed pigeon |
| HiC_scaffold_29 | 2019295 | micro | 29 | PATFA | dzscf29 | Band-tailed pigeon |
| HiC_scaffold_30 | 1962286 | micro | 30 | PATFA | dzscf30 | Band-tailed pigeon |
| HiC_scaffold_31 | 212294856 | macro | 1 | RHEAM | dzscf31 | Greater rhea |
| HiC_scaffold_2 | 161770042 | macro | 2 | RHEAM | dzscf2 | Greater rhea |
| HiC_scaffold_6 | 126853929 | macro | 3 | RHEAM | dzscf6 | Greater rhea |
| HiC_scaffold_3 | 80688299 | macro | 4 | RHEAM | dzscf3 | Greater rhea |
| HiC_scaffold_1 | 80041105 | macro | 9 | RHEAM | dzscf1 | Greater rhea |
| HiC_scaffold_30 | 69224556 | macro | 5 | RHEAM | dzscf30 | Greater rhea |
| HiC_scaffold_5 | 43122669 | macro | 7 | RHEAM | dzscf5 | Greater rhea |
| HiC_scaffold_29 | 40122881 | macro | 6 | RHEAM | dzscf29 | Greater rhea |
| HiC_scaffold_27 | 35777221 | macro | 8 | RHEAM | dzscf27 | Greater rhea |
| HiC_scaffold_25 | 29910934 | micro | 11 | RHEAM | dzscf25 | Greater rhea |
| HiC_scaffold_26 | 24624033 | micro | 12 | RHEAM | dzscf26 | Greater rhea |
| HiC_scaffold_24 | 24596671 | micro | 14 | RHEAM | dzscf24 | Greater rhea |
| HiC_scaffold_23 | 24167449 | micro | 10 | RHEAM | dzscf23 | Greater rhea |
| HiC_scaffold_28 | 23455514 | micro | 13 | RHEAM | dzscf28 | Greater rhea |
| HiC_scaffold_7 | 22135346 | micro | 15 | RHEAM | dzscf7 | Greater rhea |
| HiC_scaffold_9 | 19624805 | micro | 16 | RHEAM | dzscf9 | Greater rhea |
| HiC_scaffold_22 | 17693164 | micro | 21 | RHEAM | dzscf22 | Greater rhea |
| HiC_scaffold_8 | 16882234 | micro | 17 | RHEAM | dzscf8 | Greater rhea |
| HiC_scaffold_19 | 13802169 | micro | 19 | RHEAM | dzscf19 | Greater rhea |
| HiC_scaffold_18 | 13128898 | micro | 18 | RHEAM | dzscf18 | Greater rhea |
| HiC_scaffold_10 | 12812575 | micro | 20 | RHEAM | dzscf10 | Greater rhea |
| HiC_scaffold_17 | 9000849 | micro | 22 | RHEAM | dzscf17 | Greater rhea |
| HiC_scaffold_16 | 8268599 | micro | 24 | RHEAM | dzscf16 | Greater rhea |
| HiC_scaffold_20 | 8195719 | micro | 25 | RHEAM | dzscf20 | Greater rhea |
| HiC_scaffold_4 | 7452025 | micro | 27 | RHEAM | dzscf4 | Greater rhea |
| HiC_scaffold_15 | 6498333 | micro | 28 | RHEAM | dzscf15 | Greater rhea |
| HiC_scaffold_14 | 6044554 | micro | 29 | RHEAM | dzscf14 | Greater rhea |
| HiC_scaffold_21 | 5463505 | micro | 23 | RHEAM | dzscf21 | Greater rhea |
| HiC_scaffold_12 | 3193995 | micro | 26 | RHEAM | dzscf12 | Greater rhea |
| HiC_scaffold_13 | 2850818 | micro | 30 | RHEAM | dzscf13 | Greater rhea |
| HiC_scaffold_1 | 353375234 | macro | 1 | INTLE | dzscf1 | Eastern water dragon |
| HiC_scaffold_2 | 310803808 | macro | 2 | INTLE | dzscf2 | Eastern water dragon |
| HiC_scaffold_3 | 268945440 | macro | 4 | INTLE | dzscf3 | Eastern water dragon |
| HiC_scaffold_4 | 243750682 | macro | 3 | INTLE | dzscf4 | Eastern water dragon |
| HiC_scaffold_5 | 199434357 | macro | 5 | INTLE | dzscf5 | Eastern water dragon |
| HiC_scaffold_6 | 127569639 | macro | 6 | INTLE | dzscf6 | Eastern water dragon |
| HiC_scaffold_7 | 31587450 | micro | 8 | INTLE | dzscf7 | Eastern water dragon |
| HiC_scaffold_8 | 29318704 | micro | 10 | INTLE | dzscf8 | Eastern water dragon |
| HiC_scaffold_9 | 29106120 | micro | 7 | INTLE | dzscf9 | Eastern water dragon |
| HiC_scaffold_10 | 28455099 | micro | 9 | INTLE | dzscf10 | Eastern water dragon |
| HiC_scaffold_11 | 27099500 | micro | 11 | INTLE | dzscf11 | Eastern water dragon |
| HiC_scaffold_12 | 23275578 | micro | 12 | INTLE | dzscf12 | Eastern water dragon |
| HiC_scaffold_13 | 22219463 | micro | 13 | INTLE | dzscf13 | Eastern water dragon |
| HiC_scaffold_14 | 21106464 | micro | 14 | INTLE | dzscf14 | Eastern water dragon |
| HiC_scaffold_15 | 13809248 | micro | 20 | INTLE | dzscf15 | Eastern water dragon |
| HiC_scaffold_16 | 13196741 | micro | 21 | INTLE | dzscf16 | Eastern water dragon |
| HiC_scaffold_17 | 12608347 | micro | 15 | INTLE | dzscf17 | Eastern water dragon |
| HiC_scaffold_18 | 12072054 | micro | 22 | INTLE | dzscf18 | Eastern water dragon |
| HiC_scaffold_19 | 6800920 | micro | 16 | INTLE | dzscf19 | Eastern water dragon |
| HiC_scaffold_20 | 6465513 | micro | 23 | INTLE | dzscf20 | Eastern water dragon |
| HiC_scaffold_21 | 3006139 | micro | 17 | INTLE | dzscf21 | Eastern water dragon |
| HiC_scaffold_22 | 2224500 | micro | 18 | INTLE | dzscf22 | Eastern water dragon |
| HiC_scaffold_23 | 1625264 | micro | 19 | INTLE | dzscf23 | Eastern water dragon |
| HiC_scaffold_24 | 1305843 | micro | 24 | INTLE | dzscf24 | Eastern water dragon |
| HiC_scaffold_25 | 1046080 | micro | 25 | INTLE | dzscf25 | Eastern water dragon |
| NC_006088.5 | 197608386 | macro | 1 | CHICK | chr1 | Chicken |
| NC_006089.5 | 149682049 | macro | 2 | CHICK | chr2 | Chicken |
| NC_006090.5 | 110838418 | macro | 3 | CHICK | chr3 | Chicken |
| NC_006091.5 | 91315245 | macro | 4 | CHICK | chr4 | Chicken |
| NC_006092.5 | 59809098 | macro | 5 | CHICK | chr5 | Chicken |
| NC_006093.5 | 36374701 | macro | 6 | CHICK | chr6 | Chicken |
| NC_006094.5 | 36742308 | macro | 7 | CHICK | chr7 | Chicken |
| NC_006095.5 | 30219446 | macro | 8 | CHICK | chr8 | Chicken |
| NC_006096.5 | 24153086 | micro | 10 | CHICK | chr9 | Chicken |
| NC_006097.5 | 21119840 | micro | 11 | CHICK | chr10 | Chicken |
| NC_006098.5 | 20200042 | micro | 12 | CHICK | chr11 | Chicken |
| NC_006099.5 | 20387278 | micro | 13 | CHICK | chr12 | Chicken |
| NC_006100.5 | 19166714 | micro | 14 | CHICK | chr13 | Chicken |
| NC_006101.5 | 16219308 | micro | 15 | CHICK | chr14 | Chicken |
| NC_006102.5 | 13062184 | micro | 16 | CHICK | chr15 | Chicken |
| NC_006103.5 | 2844601 | micro | 17 | CHICK | chr16 | Chicken |
| NC_006104.5 | 10762512 | micro | 18 | CHICK | chr17 | Chicken |
| NC_006105.5 | 11373140 | micro | 19 | CHICK | chr18 | Chicken |
| NC_006106.5 | 10323212 | micro | 20 | CHICK | chr19 | Chicken |
| NC_006107.5 | 13897287 | micro | 21 | CHICK | chr20 | Chicken |
| NC_006108.5 | 6844979 | micro | 22 | CHICK | chr21 | Chicken |
| NC_006109.5 | 5459462 | micro | 23 | CHICK | chr22 | Chicken |
| NC_006110.5 | 6149580 | micro | 24 | CHICK | chr23 | Chicken |
| NC_006111.5 | 6491222 | micro | 25 | CHICK | chr24 | Chicken |
| NC_006112.4 | 3980610 | micro | 26 | CHICK | chr25 | Chicken |
| NC_006113.5 | 6055710 | micro | 27 | CHICK | chr26 | Chicken |
| NC_006114.5 | 8080432 | micro | 28 | CHICK | chr27 | Chicken |
| NC_006115.5 | 5116882 | micro | 29 | CHICK | chr28 | Chicken |
| NC_028739.2 | 1818525 | micro | 30 | CHICK | chr30 | Chicken |
| NC_028740.2 | 6153034 | micro | 31 | CHICK | chr31 | Chicken |
| NC_006119.4 | 725831 | micro | 32 | CHICK | chr32 | Chicken |
| NC_008465.4 | 7821666 | micro | 33 | CHICK | chr33 | Chicken |
| NC_006126.5 | 6813114 | macro | 34 | CHICK | chrW | Chicken |
| NC_006127.5 | 82529921 | macro | 9 | CHICK | chrZ | Chicken |
| NC_048298.1 | 347660827 | macro | 1 | TRASE | chr1 | Red-eared slider turtle |
| NC_048299.1 | 283098403 | macro | 2 | TRASE | chr2 | Red-eared slider turtle |
| NC_048300.1 | 201239519 | macro | 3 | TRASE | chr3 | Red-eared slider turtle |
| NC_048301.1 | 142343550 | macro | 5 | TRASE | chr4 | Red-eared slider turtle |
| NC_048302.1 | 140411086 | macro | 4 | TRASE | chr5 | Red-eared slider turtle |
| NC_048303.1 | 129675691 | macro | 9 | TRASE | chr6 | Red-eared slider turtle |
| NC_048304.1 | 126808733 | macro | 6 | TRASE | chr7 | Red-eared slider turtle |
| NC_048305.1 | 109307977 | macro | 8 | TRASE | chr8 | Red-eared slider turtle |
| NC_048306.1 | 104972506 | macro | 11 | TRASE | chr9 | Red-eared slider turtle |
| NC_048307.1 | 85829911 | macro | 10 | TRASE | chr10 | Red-eared slider turtle |
| NC_048308.1 | 76047897 | macro | 7 | TRASE | chr11 | Red-eared slider turtle |
| NC_048309.1 | 45061502 | micro | 17 | TRASE | chr12 | Red-eared slider turtle |
| NC_048310.1 | 43716676 | micro | 12 | TRASE | chr13 | Red-eared slider turtle |
| NC_048311.1 | 42221082 | micro | 15 | TRASE | chr14 | Red-eared slider turtle |
| NC_048312.1 | 32210117 | micro | 13 | TRASE | chr15 | Red-eared slider turtle |
| NC_048313.1 | 30045477 | micro | 24 | TRASE | chr16 | Red-eared slider turtle |
| NC_048314.1 | 26830997 | micro | 14 | TRASE | chr17 | Red-eared slider turtle |
| NC_048315.1 | 24772400 | micro | 16 | TRASE | chr18 | Red-eared slider turtle |
| NC_048316.1 | 22813715 | micro | 18 | TRASE | chr19 | Red-eared slider turtle |
| NC_048317.1 | 19368400 | micro | 19 | TRASE | chr20 | Red-eared slider turtle |
| NC_048318.1 | 19049219 | micro | 20 | TRASE | chr21 | Red-eared slider turtle |
| NC_048319.1 | 16439607 | micro | 23 | TRASE | chr22 | Red-eared slider turtle |
| NC_048320.1 | 16074632 | micro | 22 | TRASE | chr23 | Red-eared slider turtle |
| NC_048321.1 | 15148950 | micro | 21 | TRASE | chr24 | Red-eared slider turtle |
| NC_048322.1 | 7130772 | micro | 25 | TRASE | chr25 | Red-eared slider turtle |
| CM023912.1 | 117486409 | macro | 2 | CORMO | chr1 | Jackdaw |
| CM023913.1 | 154487646 | macro | 3 | CORMO | chr2 | Jackdaw |
| CM023914.1 | 110910713 | macro | 4 | CORMO | chr3 | Jackdaw |
| CM023915.1 | 73157438 | macro | 5 | CORMO | chr4 | Jackdaw |
| CM023916.1 | 64157287 | macro | 6 | CORMO | chr5 | Jackdaw |
| CM023917.1 | 36532927 | macro | 7 | CORMO | chr6 | Jackdaw |
| CM023918.1 | 38277055 | macro | 8 | CORMO | chr7 | Jackdaw |
| CM023919.1 | 32060782 | macro | 9 | CORMO | chr8 | Jackdaw |
| CM023920.1 | 26026359 | micro | 12 | CORMO | chr9 | Jackdaw |
| CM023921.1 | 20701202 | micro | 13 | CORMO | chr10 | Jackdaw |
| CM023922.1 | 20977607 | micro | 14 | CORMO | chr11 | Jackdaw |
| CM023923.1 | 21032122 | micro | 15 | CORMO | chr12 | Jackdaw |
| CM023924.1 | 19121564 | micro | 16 | CORMO | chr13 | Jackdaw |
| CM023925.1 | 16609512 | micro | 17 | CORMO | chr14 | Jackdaw |
| CM023926.1 | 11237182 | micro | 18 | CORMO | chr15 | Jackdaw |
| CM023927.1 | 11032243 | micro | 19 | CORMO | chr17 | Jackdaw |
| CM023928.1 | 11176533 | micro | 20 | CORMO | chr18 | Jackdaw |
| CM023929.1 | 11217504 | micro | 21 | CORMO | chr19 | Jackdaw |
| CM023930.1 | 14997001 | micro | 22 | CORMO | chr20 | Jackdaw |
| CM023931.1 | 6847630 | micro | 23 | CORMO | chr21 | Jackdaw |
| CM023932.1 | 4839898 | micro | 24 | CORMO | chr22 | Jackdaw |
| CM023933.1 | 8585400 | micro | 25 | CORMO | chr23 | Jackdaw |
| CM023934.1 | 6907201 | micro | 26 | CORMO | chr24 | Jackdaw |
| CM023935.1 | 6615938 | micro | 27 | CORMO | chr26 | Jackdaw |
| CM023936.1 | 4573168 | micro | 28 | CORMO | chr27 | Jackdaw |
| CM023937.1 | 5735232 | micro | 29 | CORMO | chr28 | Jackdaw |
| CM023938.1 | 79982351 | macro | 10 | CORMO | chrZ | Jackdaw |
| CM023939.1 | 73777161 | macro | 1 | CORMO | chr1A | Jackdaw |
| CM023940.1 | 20605299 | micro | 11 | CORMO | chr4A | Jackdaw |
| CM019148.1 | 375026955 | macro | 1 | NAJNA | chr1 | Indian cobra |
| CM019149.1 | 309871788 | macro | 2 | NAJNA | chr2 | Indian cobra |
| CM019150.1 | 224088900 | macro | 3 | NAJNA | chr3 | Indian cobra |
| CM019151.1 | 191024962 | macro | 6 | NAJNA | chr4 | Indian cobra |
| CM019152.1 | 62775154 | macro | 5 | NAJNA | chr5 | Indian cobra |
| CM019153.1 | 54008553 | macro | 4 | NAJNA | chr6 | Indian cobra |
| CM019154.1 | 53796468 | macro | 8 | NAJNA | chr7 | Indian cobra |
| CM019155.1 | 154607413 | macro | 9 | NAJNA | chrZ | Indian cobra |
| CM019156.1 | 32026552 | micro | 11 | NAJNA | chrMIC_1 | Indian cobra |
| CM019159.1 | 30264392 | micro | 7 | NAJNA | chrMIC_2 | Indian cobra |
| CM019160.1 | 29932115 | micro | 10 | NAJNA | chrMIC_3 | Indian cobra |
| CM019161.1 | 25974797 | micro | 13 | NAJNA | chrMIC_4 | Indian cobra |
| CM019162.1 | 25174854 | micro | 12 | NAJNA | chrMIC_5 | Indian cobra |
| CM019163.1 | 23367679 | micro | 14 | NAJNA | chrMIC_6 | Indian cobra |
| CM019164.1 | 20042958 | micro | 15 | NAJNA | chrMIC_7 | Indian cobra |
| CM019165.1 | 17750283 | micro | 16 | NAJNA | chrMIC_8 | Indian cobra |
| CM019166.1 | 15339066 | micro | 17 | NAJNA | chrMIC_9 | Indian cobra |
| CM019157.1 | 11847348 | micro | 18 | NAJNA | chrMIC_10 | Indian cobra |
| CM019158.1 | 10772589 | micro | 19 | NAJNA | chrMIC_11 | Indian cobra |
| NC_041728.1 | 186506016 | macro | 1 | ORNAN | chr1 | Platypus |
| NC_041729.1 | 171719663 | macro | 2 | ORNAN | chr2 | Platypus |
| NC_041730.1 | 141987082 | macro | 3 | ORNAN | chr3 | Platypus |
| NC_041731.1 | 136094389 | macro | 4 | ORNAN | chr4 | Platypus |
| NC_041732.1 | 110293030 | macro | 5 | ORNAN | chr5 | Platypus |
| NC_041733.1 | 51493492 | macro | 6 | ORNAN | chr6 | Platypus |
| NC_041734.1 | 83338043 | macro | 7 | ORNAN | chr7 | Platypus |
| NC_041735.1 | 70949814 | macro | 8 | ORNAN | chr8 | Platypus |
| NC_041736.1 | 61861223 | macro | 9 | ORNAN | chr9 | Platypus |
| NC_041737.1 | 59444696 | macro | 10 | ORNAN | chr10 | Platypus |
| NC_041738.1 | 63506031 | macro | 11 | ORNAN | chr11 | Platypus |
| NC_041739.1 | 60456049 | macro | 12 | ORNAN | chr12 | Platypus |
| NC_041740.1 | 42294642 | macro | 13 | ORNAN | chr13 | Platypus |
| NC_041741.1 | 52889406 | macro | 14 | ORNAN | chr14 | Platypus |
| NC_041742.1 | 26375019 | macro | 15 | ORNAN | chr15 | Platypus |
| NC_041743.1 | 46397275 | macro | 16 | ORNAN | chr16 | Platypus |
| NC_041744.1 | 45536670 | macro | 17 | ORNAN | chr17 | Platypus |
| NC_041745.1 | 44684501 | macro | 18 | ORNAN | chr18 | Platypus |
| NC_041746.1 | 35120541 | macro | 19 | ORNAN | chr19 | Platypus |
| NC_041747.1 | 30535534 | macro | 20 | ORNAN | chr20 | Platypus |
| NC_041748.1 | 24514859 | macro | 21 | ORNAN | chr21 | Platypus |
| NC_041749.1 | 125040914 | macro | 22 | ORNAN | chrX1 | Platypus |
| NC_041750.1 | 29662106 | macro | 23 | ORNAN | chrX2 | Platypus |
| NC_041751.1 | 33863336 | macro | 24 | ORNAN | chrX3 | Platypus |
| NC_041752.1 | 8639456 | macro | 25 | ORNAN | chrX4 | Platypus |
| NC_041753.1 | 70139320 | macro | 26 | ORNAN | chrX5 | Platypus |
| CM026898.1 | 348265484 | macro | 1 | CHEMY | chr1 | Green sea turtle |
| CM026899.1 | 262513884 | macro | 2 | CHEMY | chr2 | Green sea turtle |
| CM026900.1 | 204120564 | macro | 3 | CHEMY | chr3 | Green sea turtle |
| CM026901.1 | 142324156 | macro | 4 | CHEMY | chr4 | Green sea turtle |
| CM026902.1 | 134428053 | macro | 5 | CHEMY | chr5 | Green sea turtle |
| CM026903.1 | 133272604 | macro | 9 | CHEMY | chr6 | Green sea turtle |
| CM026904.1 | 123872898 | macro | 6 | CHEMY | chr7 | Green sea turtle |
| CM026905.1 | 106505537 | macro | 8 | CHEMY | chr8 | Green sea turtle |
| CM026906.1 | 101620591 | macro | 11 | CHEMY | chr9 | Green sea turtle |
| CM026907.1 | 84054250 | macro | 10 | CHEMY | chr10 | Green sea turtle |
| CM026908.1 | 79522289 | macro | 7 | CHEMY | chr11 | Green sea turtle |
| CM026909.1 | 44456234 | micro | 13 | CHEMY | chr12 | Green sea turtle |
| CM026910.1 | 42953438 | micro | 14 | CHEMY | chr13 | Green sea turtle |
| CM026911.1 | 42626581 | micro | 12 | CHEMY | chr14 | Green sea turtle |
| CM026912.1 | 33324608 | micro | 15 | CHEMY | chr15 | Green sea turtle |
| CM026913.1 | 25690524 | micro | 16 | CHEMY | chr16 | Green sea turtle |
| CM026914.1 | 25202864 | micro | 17 | CHEMY | chr17 | Green sea turtle |
| CM026915.1 | 23386534 | micro | 18 | CHEMY | chr18 | Green sea turtle |
| CM026916.1 | 19769559 | micro | 19 | CHEMY | chr19 | Green sea turtle |
| CM026917.1 | 19256569 | micro | 20 | CHEMY | chr20 | Green sea turtle |
| CM026918.1 | 19202859 | micro | 21 | CHEMY | chr21 | Green sea turtle |
| CM026919.1 | 11152253 | micro | 23 | CHEMY | chr22 | Green sea turtle |
| CM026920.1 | 9977174 | micro | 22 | CHEMY | chr23 | Green sea turtle |
| CM026921.1 | 18854909 | micro | 24 | CHEMY | chr24 | Green sea turtle |
| CM026922.1 | 16433863 | micro | 25 | CHEMY | chr25 | Green sea turtle |
| CM026923.1 | 16225734 | micro | 26 | CHEMY | chr26 | Green sea turtle |
| CM026924.1 | 16135183 | micro | 27 | CHEMY | chr27 | Green sea turtle |
| CM026925.1 | 7864902 | micro | 28 | CHEMY | chr28 | Green sea turtle |
| NC_041312.1 | 140209921 | macro | 2 | PODMU | chr1 | Common wall lizard |
| NC_041313.1 | 128038163 | macro | 4 | PODMU | chr2 | Common wall lizard |
| NC_041314.1 | 124954059 | macro | 1 | PODMU | chr3 | Common wall lizard |
| NC_041315.1 | 107526675 | macro | 7 | PODMU | chr4 | Common wall lizard |
| NC_041316.1 | 100393251 | macro | 8 | PODMU | chr5 | Common wall lizard |
| NC_041317.1 | 99678440 | macro | 6 | PODMU | chr6 | Common wall lizard |
| NC_041318.1 | 92398148 | macro | 5 | PODMU | chr7 | Common wall lizard |
| NC_041319.1 | 90342841 | macro | 11 | PODMU | chr8 | Common wall lizard |
| NC_041320.1 | 79135106 | macro | 9 | PODMU | chr9 | Common wall lizard |
| NC_041321.1 | 76280819 | macro | 10 | PODMU | chr10 | Common wall lizard |
| NC_041322.1 | 65231656 | macro | 3 | PODMU | chr11 | Common wall lizard |
| NC_041323.1 | 60949594 | macro | 12 | PODMU | chr12 | Common wall lizard |
| NC_041324.1 | 56546671 | macro | 13 | PODMU | chr13 | Common wall lizard |
| NC_041325.1 | 54988359 | macro | 14 | PODMU | chr14 | Common wall lizard |
| NC_041326.1 | 46441056 | micro | 16 | PODMU | chr15 | Common wall lizard |
| NC_041327.1 | 43659005 | micro | 17 | PODMU | chr16 | Common wall lizard |
| NC_041328.1 | 42925489 | micro | 18 | PODMU | chr17 | Common wall lizard |
| NC_041329.1 | 14237138 | micro | 19 | PODMU | chr18 | Common wall lizard |
| NC_041330.1 | 50871175 | macro | 15 | PODMU | chrZ | Common wall lizard |
| NC_048605.1 | 131769443 | macro | 2 | ZOOVI | LG1 | Common lizard |
| NC_048606.1 | 117834363 | macro | 5 | ZOOVI | LG2 | Common lizard |
| NC_048607.1 | 116371480 | macro | 1 | ZOOVI | LG3 | Common lizard |
| NC_048608.1 | 100631306 | macro | 11 | ZOOVI | LG4 | Common lizard |
| NC_048609.1 | 98023110 | macro | 8 | ZOOVI | LG5 | Common lizard |
| NC_048610.1 | 94199661 | macro | 13 | ZOOVI | LG6 | Common lizard |
| NC_048611.1 | 92810032 | macro | 7 | ZOOVI | LG7 | Common lizard |
| NC_048612.1 | 79626714 | macro | 6 | ZOOVI | LG8 | Common lizard |
| NC_048613.1 | 69679922 | macro | 12 | ZOOVI | LG9 | Common lizard |
| NC_048614.1 | 54413957 | macro | 14 | ZOOVI | LG10 | Common lizard |
| NC_048615.1 | 51059636 | macro | 16 | ZOOVI | LG11 | Common lizard |
| NC_048616.1 | 49589567 | micro | 10 | ZOOVI | LG12 | Common lizard |
| NC_048617.1 | 48213930 | micro | 15 | ZOOVI | LG13 | Common lizard |
| NC_048618.1 | 46006402 | micro | 9 | ZOOVI | LG14 | Common lizard |
| NC_048619.1 | 44540502 | micro | 17 | ZOOVI | LG15 | Common lizard |
| NC_048620.1 | 40703745 | micro | 19 | ZOOVI | LG16 | Common lizard |
| NC_048621.1 | 39436030 | micro | 18 | ZOOVI | LG17 | Common lizard |
| NC_048622.1 | 35462695 | micro | 4 | ZOOVI | LG18 | Common lizard |
| NC_048623.1 | 24898947 | micro | 3 | ZOOVI | LG19 | Common lizard |
| CM012306.1 | 311712589 | macro | 1 | CROVV | chr1 | Prairie rattlesnake |
| CM012307.1 | 230514893 | macro | 2 | CROVV | chr2 | Prairie rattlesnake |
| CM012308.1 | 179897795 | macro | 3 | CROVV | chr3 | Prairie rattlesnake |
| CM012309.1 | 103290658 | macro | 4 | CROVV | chr4 | Prairie rattlesnake |
| CM012310.1 | 87326383 | macro | 5 | CROVV | chr5 | Prairie rattlesnake |
| CM012311.1 | 69529916 | macro | 7 | CROVV | chr6 | Prairie rattlesnake |
| CM012312.1 | 61662597 | macro | 6 | CROVV | chr7 | Prairie rattlesnake |
| CM012313.1 | 22521304 | micro | 10 | CROVV | chr9 | Prairie rattlesnake |
| CM012314.1 | 19978502 | micro | 9 | CROVV | chr10 | Prairie rattlesnake |
| CM012315.1 | 16785021 | micro | 14 | CROVV | chr11 | Prairie rattlesnake |
| CM012316.1 | 16736725 | micro | 11 | CROVV | chr12 | Prairie rattlesnake |
| CM012317.1 | 13784772 | micro | 15 | CROVV | chr13 | Prairie rattlesnake |
| CM012318.1 | 13760810 | micro | 12 | CROVV | chr14 | Prairie rattlesnake |
| CM012319.1 | 12380204 | micro | 18 | CROVV | chr15 | Prairie rattlesnake |
| CM012320.1 | 10971997 | micro | 13 | CROVV | chr16 | Prairie rattlesnake |
| CM012321.1 | 6767157 | micro | 16 | CROVV | chr17 | Prairie rattlesnake |
| CM012322.1 | 5372825 | micro | 17 | CROVV | chr18 | Prairie rattlesnake |
| CM012323.1 | 113984505 | macro | 8 | CROVV | chrZ | Prairie rattlesnake |
| NC_044244.1 | 197551010 | macro | 1 | CALAN | chr1 | Anna's hummingbird |
| NC_044245.1 | 151342139 | macro | 2 | CALAN | chr2 | Anna's hummingbird |
| NC_044246.1 | 114810999 | macro | 3 | CALAN | chr3 | Anna's hummingbird |
| NC_044247.1 | 18597117 | micro | 12 | CALAN | chr4 | Anna's hummingbird |
| NC_044248.1 | 44745344 | macro | 5 | CALAN | chr4A | Anna's hummingbird |
| NC_044249.1 | 23291945 | micro | 4 | CALAN | chr4B | Anna's hummingbird |
| NC_044250.1 | 16645885 | micro | 7 | CALAN | chr5 | Anna's hummingbird |
| NC_044251.1 | 43880846 | macro | 6 | CALAN | chr5A | Anna's hummingbird |
| NC_044252.1 | 35401958 | macro | 8 | CALAN | chr6 | Anna's hummingbird |
| NC_044253.1 | 39139214 | macro | 9 | CALAN | chr7 | Anna's hummingbird |
| NC_044254.1 | 31090148 | macro | 10 | CALAN | chr8 | Anna's hummingbird |
| NC_044255.1 | 25686456 | micro | 13 | CALAN | chr9 | Anna's hummingbird |
| NC_044256.1 | 22664390 | micro | 14 | CALAN | chr10 | Anna's hummingbird |
| NC_044257.1 | 20302349 | micro | 15 | CALAN | chr11 | Anna's hummingbird |
| NC_044258.1 | 21352500 | micro | 16 | CALAN | chr12 | Anna's hummingbird |
| NC_044259.1 | 17696115 | micro | 17 | CALAN | chr13 | Anna's hummingbird |
| NC_044260.1 | 15497061 | micro | 18 | CALAN | chr14 | Anna's hummingbird |
| NC_044261.1 | 13887164 | micro | 19 | CALAN | chr15 | Anna's hummingbird |
| NC_044262.1 | 10473655 | micro | 20 | CALAN | chr17 | Anna's hummingbird |
| NC_044263.1 | 11617720 | micro | 21 | CALAN | chr18 | Anna's hummingbird |
| NC_044264.1 | 11166003 | micro | 22 | CALAN | chr19 | Anna's hummingbird |
| NC_044265.1 | 15116875 | micro | 23 | CALAN | chr20 | Anna's hummingbird |
| NC_044266.1 | 7710055 | micro | 24 | CALAN | chr21 | Anna's hummingbird |
| NC_044267.1 | 5198135 | micro | 25 | CALAN | chr22 | Anna's hummingbird |
| NC_044268.1 | 6423903 | micro | 26 | CALAN | chr23 | Anna's hummingbird |
| NC_044269.1 | 6461084 | micro | 27 | CALAN | chr24 | Anna's hummingbird |
| NC_044270.1 | 2161911 | micro | 28 | CALAN | chr25 | Anna's hummingbird |
| NC_044271.1 | 6306732 | micro | 29 | CALAN | chr26 | Anna's hummingbird |
| NC_044272.1 | 5749075 | micro | 30 | CALAN | chr27 | Anna's hummingbird |
| NC_044273.1 | 5713987 | micro | 31 | CALAN | chr28 | Anna's hummingbird |
| NC_044274.1 | 74081004 | macro | 11 | CALAN | chrZ | Anna's hummingbird |
| NC_044276.1 | 26448777 | macro | 33 | CALAN | chrW | Anna's hummingbird |
| NC_044275.1 | 1888140 | micro | 32 | CALAN | chr33 | Anna's hummingbird |
| chr1 | 248956422 | macro | 1 | HUMAN | chr1 | Human |
| chr2 | 242193529 | macro | 2 | HUMAN | chr2 | Human |
| chr3 | 198295559 | macro | 3 | HUMAN | chr3 | Human |
| chr4 | 190214555 | macro | 4 | HUMAN | chr4 | Human |
| chr5 | 181538259 | macro | 5 | HUMAN | chr5 | Human |
| chr6 | 170805979 | macro | 6 | HUMAN | chr6 | Human |
| chr7 | 159345973 | macro | 7 | HUMAN | chr7 | Human |
| chr8 | 145138636 | macro | 8 | HUMAN | chr8 | Human |
| chr9 | 138394717 | macro | 9 | HUMAN | chr9 | Human |
| chr10 | 133797422 | macro | 10 | HUMAN | chr10 | Human |
| chr11 | 135086622 | macro | 11 | HUMAN | chr11 | Human |
| chr12 | 133275309 | macro | 12 | HUMAN | chr12 | Human |
| chr13 | 114364328 | macro | 13 | HUMAN | chr13 | Human |
| chr14 | 107043718 | macro | 14 | HUMAN | chr14 | Human |
| chr15 | 101991189 | macro | 15 | HUMAN | chr15 | Human |
| chr16 | 90338345 | macro | 16 | HUMAN | chr16 | Human |
| chr17 | 83257441 | macro | 17 | HUMAN | chr17 | Human |
| chr18 | 80373285 | macro | 18 | HUMAN | chr18 | Human |
| chr19 | 58617616 | macro | 19 | HUMAN | chr19 | Human |
| chr20 | 64444167 | macro | 20 | HUMAN | chr20 | Human |
| chr21 | 46709983 | macro | 21 | HUMAN | chr21 | Human |
| chr22 | 50818468 | macro | 22 | HUMAN | chr22 | Human |
| chrX | 156040895 | macro | 23 | HUMAN | chrX | Human |
| chrY | 57227415 | macro | 24 | HUMAN | chrY | Human |
| NC_049979.1 | 35335272 | micro | 1 | BRAFL | chr1 | Florida lancelet |
| NC_049980.1 | 35136500 | micro | 2 | BRAFL | chr2 | Florida lancelet |
| NC_049981.1 | 34247348 | micro | 3 | BRAFL | chr3 | Florida lancelet |
| NC_049982.1 | 34244393 | micro | 4 | BRAFL | chr4 | Florida lancelet |
| NC_049983.1 | 29550017 | micro | 5 | BRAFL | chr5 | Florida lancelet |
| NC_049984.1 | 28086088 | micro | 6 | BRAFL | chr6 | Florida lancelet |
| NC_049985.1 | 25954402 | micro | 7 | BRAFL | chr7 | Florida lancelet |
| NC_049986.1 | 25575982 | micro | 8 | BRAFL | chr8 | Florida lancelet |
| NC_049987.1 | 25441410 | micro | 9 | BRAFL | chr9 | Florida lancelet |
| NC_049988.1 | 23931428 | micro | 10 | BRAFL | chr10 | Florida lancelet |
| NC_049989.1 | 23529748 | micro | 11 | BRAFL | chr11 | Florida lancelet |
| NC_049990.1 | 22973701 | micro | 12 | BRAFL | chr12 | Florida lancelet |
| NC_049991.1 | 22407120 | micro | 13 | BRAFL | chr13 | Florida lancelet |
| NC_049992.1 | 21813864 | micro | 14 | BRAFL | chr14 | Florida lancelet |
| NC_049993.1 | 21514203 | micro | 15 | BRAFL | chr15 | Florida lancelet |
| NC_049994.1 | 19152309 | micro | 16 | BRAFL | chr16 | Florida lancelet |
| NC_049995.1 | 19122194 | micro | 17 | BRAFL | chr17 | Florida lancelet |
| NC_049996.1 | 18554083 | micro | 18 | BRAFL | chr18 | Florida lancelet |
| NC_049997.1 | 17116508 | micro | 19 | BRAFL | chr19 | Florida lancelet |

**SI References**

1.	M. Vujošević, J. Blagojević, The distribution of constitutive heterochromatin and nucleolus organizers in lizards of the family Lacertidae (Sauria). . *Genetika* **31**, 269–276 (1999).

2.	G. Odierna *et al.*, Progressive differentiation of the W sex‐chromosome between oviparous and viviparous populations of Zootoca vivipara (Reptilia, Lacertidae). *Italian Journal of Zoology* **65**, 295-302 (1998).

3.	M. J. da Silva *et al.*, The Karyotype of Salvator merianae (Squamata, Teiidae): Analyses by Classical and Molecular Cytogenetic Techniques. *Cytogenet Genome Res* **160**, 94-99 (2020).

4.	K. Matsubara *et al.*, Molecular cloning and characterization of satellite DNA sequences from constitutive heterochromatin of the habu snake (Protobothrops flavoviridis, Viperidae) and the Burmese python (Python bivittatus, Pythonidae). *Chromosoma* **124**, 529-539 (2015).

5.	R. J. Baker, G. A. Mengden, J. J. Bull, Karyotypic Studies of Thirty-Eight Species of North American Snakes. *Copeia* **1972**, 257-265 (1972).

6.	L. Singh, Evolution of karyotypes in snakes. *Chromosoma* **38**, 185-236 (1972).

7.	H. A. Ansari, N. Takagi, M. Sasaki, Morphological differentiation of sex chromosomes in three species of ratite birds. *Cytogenetic and Genome Research* **47**, 185-188 (1988).

8.	M. Guttenbach *et al.*, Comparative chromosome painting of chicken autosomal paints 1-9 in nine different bird species. *Cytogenet Genome Res* **103**, 173-184 (2003).

9.	M. A. C. Mendonça, C. R. Carvalho, W. R. Clarindo, DNA amount of chicken chromosomes resolved by image cytometry. *Caryologia* **69**, 201-206 (2016).

10.	A. Rhie *et al.*, Towards complete and error-free genome assemblies of all vertebrate species. *Nature* **592**, 737-746 (2021).

11.	V. Jovanovic, L. Atkins, Karyotypes of four passerine birds belonging to the families Turdidae, Mimidae, and Corvidae. *Chromosoma* **26**, 388-394 (1969).

12.	R. Kretschmer *et al.*, Repetitive DNAs and shrink genomes: A chromosomal analysis in nine Columbidae species (Aves, Columbiformes). *Genet Mol Biol* **41**, 98-106 (2018).

13.	A. K. Patnaik, M. Samanta, R. Prasad, Chromosome complement and banding patterns in a pelecaniform bird, Phalacrocorax niger. *Journal of Heredity* **72**, 447-449 (1981).

14.	R. Masuda *et al.*, Genetic characteristics of endangered Japanese golden eagles (Aquila chrysaetos japonica) based on mitochondrial DNA D-loop sequences and karyotypes. *Zoo Biology* **17**, 111-121 (1998).

15.	E. M. Valleley, C. J. Harrison, Y. Cook, M. W. Ferguson, P. T. Sharpe, The karyotype of Alligator mississippiensis, and chromosomal mapping of the ZFY/X homologue, Zfc. *Chromosoma* **103**, 502-507 (1994).

16.	C. R. D. Machado *et al.*, Comparative Cytogenetics of Four Sea Turtle Species (Cheloniidae): G-Banding Pattern and in situ Localization of Repetitive DNA Units. *Cytogenet Genome Res* **160**, 531-538 (2020).

17.	F. Cleiton, L. Giuliano-Caetano, Cytogenetic characterization of two turtle species: Trachemys dorbigni and Trachemys scripta elegans. *Caryologia* **61**, 253-257 (2008).

18.	W. Rens *et al.*, The multiple sex chromosomes of platypus and echidna are not completely identical and several share homology with the avian Z. *Genome Biol* **8**, R243 (2007).

19.	J. H. Tjio, A. Levan, THE CHROMOSOME NUMBER OF MAN. *Hereditas* **42**, 1-6 (1956).

20.	T. C. Hsu, K. Benirschke, "Phascolarctos cinereus (Koala)" in An Atlas of Mammalian Chromosomes. (Springer, New York, NY, 1977), <https://doi.org/10.1007/978-1-4615-6436-2_2>.
